# A Compendium of Kinetic Cell Death Modulatory Profiles Identifies Ferroptosis Regulators

**DOI:** 10.1101/826925

**Authors:** Megan Conlon, Carson Poltorack, Giovanni C. Forcina, Alex Wells, Melodie Mallais, Alexis Kahanu, Leslie Magtanong, Derek A. Pratt, Scott J. Dixon

## Abstract

Cell death can be executed by regulated apoptotic and non-apoptotic pathways, including the iron-dependent process of ferroptosis. Small molecules are essential tools for studying the regulation of cell death. Using live-cell, time-lapse imaging, and a library of 1,833 small molecules including FDA-approved drugs and investigational agents, we assemble a large compendium of kinetic cell death ‘modulatory profiles’ for inducers of apoptosis and ferroptosis. From this dataset we identified dozens of small molecule inhibitors of ferroptosis, including numerous investigational and FDA-approved drugs with unexpected off-target antioxidant or iron chelating activities. ATP-competitive mechanistic target of rapamycin (mTOR) inhibitors, by contrast, were on-target ferroptosis inhibitors. Further investigation revealed both mTOR-dependent and mTOR-independent mechanisms linking amino acid levels to the regulation of ferroptosis sensitivity in cancer cells. These results highlight widespread bioactive compound pleiotropy and link amino acid sensing to the regulation of ferroptosis.

## Introduction

Cell death is executed by a number of distinct apoptotic and non-apoptotic cell death processes^1^. Ferroptosis is a non-apoptotic cell death process that can be triggered by reduced uptake of cystine by the system x_c_^-^ cystine/glutamate antiporter^2, 3^. Reduced cystine uptake leads to depletion of intracellular glutathione (GSH) and inactivation of the GSH-dependent phospholipid hydroperoxidase glutathione peroxidase 4 (GPX4), causing iron-dependent accumulation of toxic lipid hydroperoxides^4, 5^. Small molecules that can induce ferroptosis may be useful to selectively target certain cancer cells^4, 6, 7^. By contrast, small molecules that can inhibit this process may be useful to prevent cell death in diverse pathological contexts^8, 9^. Greater insights into the basic mechanisms of ferroptosis, along with modulators of this process suitable for clinical use, are urgently needed^10^.

Synthetic small molecules and natural products are potent perturbagens that enable powerful chemical enhancer/suppressor (“modulator”) screens to characterize cell death mechanisms and uncover cell death inhibitors to treat disease^11–14^. Apart from the intended target, small molecules often have off-target effects on other proteins or processes. While these effects can confound the interpretation of large-scale studies, careful analysis can pinpoint unexpected mechanisms of action and new drug repurposing opportunities^12, 15–18^. Most large-scale drug repurposing studies focus on protein off-targets, but compound-specific chemical reactivities that can contribute to unanticipated biological effects within the cell, independent of protein engagement, are often ignored^19^.

Towards the goal of identifying modulators of cell death and drug repurposing candidates, we generated a compendium of cell death modulatory profiles using arrayed high-throughput time-lapse population cell death imaging^20^ and a library of annotated bioactive compounds. Analysis of this compendium identified potent small molecule inhibitors of ferroptosis, including several existing FDA-approved drugs with unanticipated radical-trapping antioxidant (RTA) activities. We also pinpointed inhibitors of the mechanistic target of rapamycin (mTOR) pathway as on-target modulators of ferroptosis and dissected the role of mTOR-dependent and mTOR-independent amino acid sensing mechanisms in this process. This work establishes a new resource to understand the chemical modulation of apoptotic and non-apoptotic cell death pathways including ferroptosis.

## Results

### Large-scale imaging-based cell death modulatory profiling

Profiles of phenotypic similarity across hundreds of perturbations can be used to classify compound mechanism of action and identify novel compound activities^11, 21, 22^. We recently developed a sensitive method to directly assess population cell death, scalable time-lapse analysis of cell death kinetics (STACK)^20^. To identify novel regulators of cell death, we used STACK to analyze cell death over time in HT-1080^N^ fibrosarcoma cells treated with the pro-apoptotic agents thapsigargin, tunicamycin, camptothecin, etoposide, bortezomib or vinblastine, and the pro-ferroptotic agents erastin, sorafenib or ML162, each tested at a single lethal concentration. For each lethal query, we examined the ability of 1,833 different compounds, including FDA-approved drugs, investigational agents and natural products, to modulate cell death over time (**Fig. 1a,b**). Each compound was tested at a fixed dose of 5 µM, which in a previous study allowed for the effective identification of cell death modulators^20^. HT-1080^N^ cells were treated with query and modulator compounds and then imaged every two h for 24 to 112 h, depending on the lethal compound (**Fig. 1b**). The resultant dataset ultimately comprised ∼700,000 population measurements from a total of ∼16,000 distinct compound combinations and control conditions.

**Figure 1.**
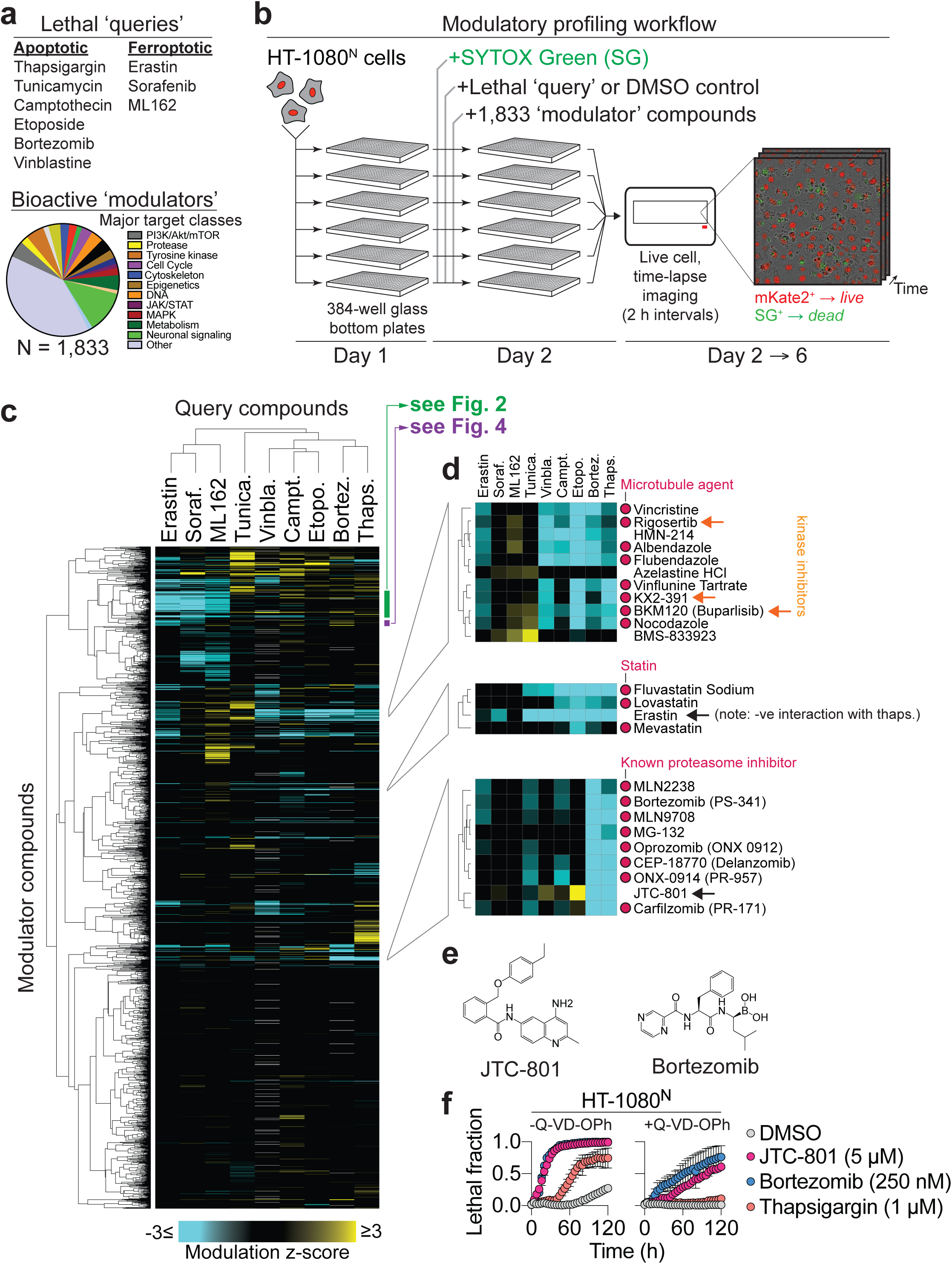
Overview of the time-lapse modulatory profiling approach. **a**, Summary of lethal query and bioactive compound modulators. **b**, Overview of compound profiling workflow. NLS::mKate2-expressing HT-1080 (i.e. HT-1080^N^) cells and small molecules were added to the wells of six 384-well plates using a robotic liquid handler. Each well from each plate was imaged every two hours to identify live cells expressing NLS::mKate2 and dead cells positive for the uptake of SYTOX Green. Counts of live and dead cells were used to determine population cell death over time and examine modulatory interactions^20^ (see also **Supplementary Fig. 1**). **c**, Heatmap summary of cell death modulation by 1,833 bioactive compounds. Clusters of interest are highlighted to the right. **d**, Expansion of heatmap sub-clusters enriched for microtubule inhibitors, ferroptosis-inducing compounds, and proteasome inhibitors. Orange arrows indicate compounds with known off-target tubulin binding. Red arrow indicates a new predicted proteasome inhibitor. **e**, Structures of JTC-801 and a known proteasome inhibitor, bortezomib. **f**, Cell death over time ± the pan-caspase inhibitor Q-VD-OPh (25 µM). Results are mean ± SD from three independent experiments.

HT-1080^N^ cells stably express the live cell marker nuclear-localized mKate2 and were incubated in media containing the dead cell marker SYTOX Green, enabling the automated identification of live and dead cells using STACK. Counts of live and dead cells were used to compute the lethal fraction over time^20^, and deviation from the expected effect of each query-modulator combination was calculated using the Bliss independence model^23, 24^ (**Supplementary Fig. 1a,b**). Deviation z-scores were hierarchically clustered in an unsupervised manner across query and modulator compounds and summarized as a heat map (**Fig. 1c**). Ferroptosis-inducing and apoptosis-inducing compounds were clearly segregated as a function of their modulation by the 1,833 bioactive modulator compounds, consistent with distinct biochemical mechanisms of action. Functionally related clusters of modulator compounds were likewise apparent, with patterns of clustering providing insight into mechanism of action under these conditions (**Fig. 1d**). For example, several microtubule destabilizing agents, including vincristine and vinflunine, clustered with the kinase inhibitors rigosertib, BKM120 (buparlisib) and KX2-391 (orange arrows in **Fig. 1d**), entirely consistent with recent reports that all three of these kinase inhibitors can also directly block tubulin polymerization^25–27^. Similarly, erastin (also present in the library as a modulator compound) clustered with three statins (fluvastatin, lovastatin and mevastatin), in line with the finding that inhibition of 3-hydroxy-3-methyl-glutaryl-coenzyme A reductase by statins can promote ferroptosis^7, 28^. Here, the clustering of erastin with statins is not due to the ability of erastin to inhibit the same target as the statins, but presumably due to similar functional effects on cell death. The opioid receptor antagonist JTC-801 clustered unexpectedly with a number of proteasome inhibitors (**Figure 1d**). This molecule was recently shown to induce apoptosis in cancer cells through a poorly understood mechanism^29, 30^. Re-testing in HT-1080^N^ cells showed that JTC-801 induced cell death with similar rapid kinetics and partial sensitivity to the pan-caspase inhibitor Q-VD-OPh as the proteasome inhibitor bortezomib, and that these effects were distinct from those caused by a different apoptosis inducer, thapsigargin (**Fig. 1e,f**). Thus, JTC-801 may inhibit the function of the proteasome or a closely related process, and our results caution against the use of this molecule as a modulator of opioid receptor function.

A notable observation within this dataset were apparent widespread negative interactions where the combination of two molecules produced less cell death than what is predicted from the lethality of each individual compound alone. For example, the combination of any compound with itself (e.g. bortezomib with bortezomib) produced less cell death than anticipated and was accordingly scored as a negative (blue) interaction (**Fig. 1d**). This is explained in part by the known mathematical limitation of the Bliss model in accounting for self-by-self drug interactions^31^. However, in other cases, the two distinct lethal compounds also produced apparent negative interactions. In some cases, the lethal effect of a rapidly-acting lethal compound could mask the effects of a more slowly-acting lethal compound, although this will require more investigation. In other instances, we speculated that two distinct lethal compounds could produce less death than expected or even inhibitory (i.e. suppressive)^32^ effects. To explore this possibility further, we examined the negative interaction observed between the lethal sarco/endoplasmic reticulum Ca^2+^ ATPase inhibitor query thapsigargin and the lethal system x_c_^-^ inhibitor erastin (**Fig. 1c,d**). In two unrelated cell lines, thapsigargin cotreatment potently suppressed the lethality of the erastin analog erastin2, but not the lethality of RSL3, consistent with the negative interaction detected in the primary screen (**Supplementary Fig. 1c**). Further exploration of this dataset may yield additional unexpected compound-compound interactions, both positive and negative.

### Cell-based characterization of new antioxidants and iron chelators

Within the compendium, our attention was attracted to a large cluster of 48 compounds enriched for potent suppressors of ferroptosis but not apoptosis (**Fig. 2a**, green bar in **Fig. 1c**). This cluster included the known radical trapping antioxidants ferrostatin-1 (Fer-1)^2, 33^ and phenothiazine^34^, as well as the MEK1/2 inhibitor U0126 which, consistent with a recent report^35^, is most likely to inhibit ferroptosis through an antioxidant effect (**Supplementary Fig. 2**). Other compounds in this cluster, such as the adrenergic receptor inhibitor carvedilol^36^, were known antioxidants but not previously shown to block ferroptosis. The remaining compounds in this cluster had diverse structures and annotated targets, such as the NF-κB pathway (JSH-23)^37^, sphingosine kinase (SKI II)^38^, and vascular endothelial growth factor receptor (linifanib)^39^. Structurally, many of these compounds contained conjugated primary or secondary amines reminiscent of the structure of ferrostatin-1 (**Fig. 2b**). Several other compounds were clear outliers in their ability to inhibit ferroptosis compared to other compounds with the same annotated protein target, hinting at likely off-target activities (**Supplementary Fig. 3a**). We hypothesized that the compounds in this cluster inhibited ferroptosis via antioxidant effects.

**Figure 2.**
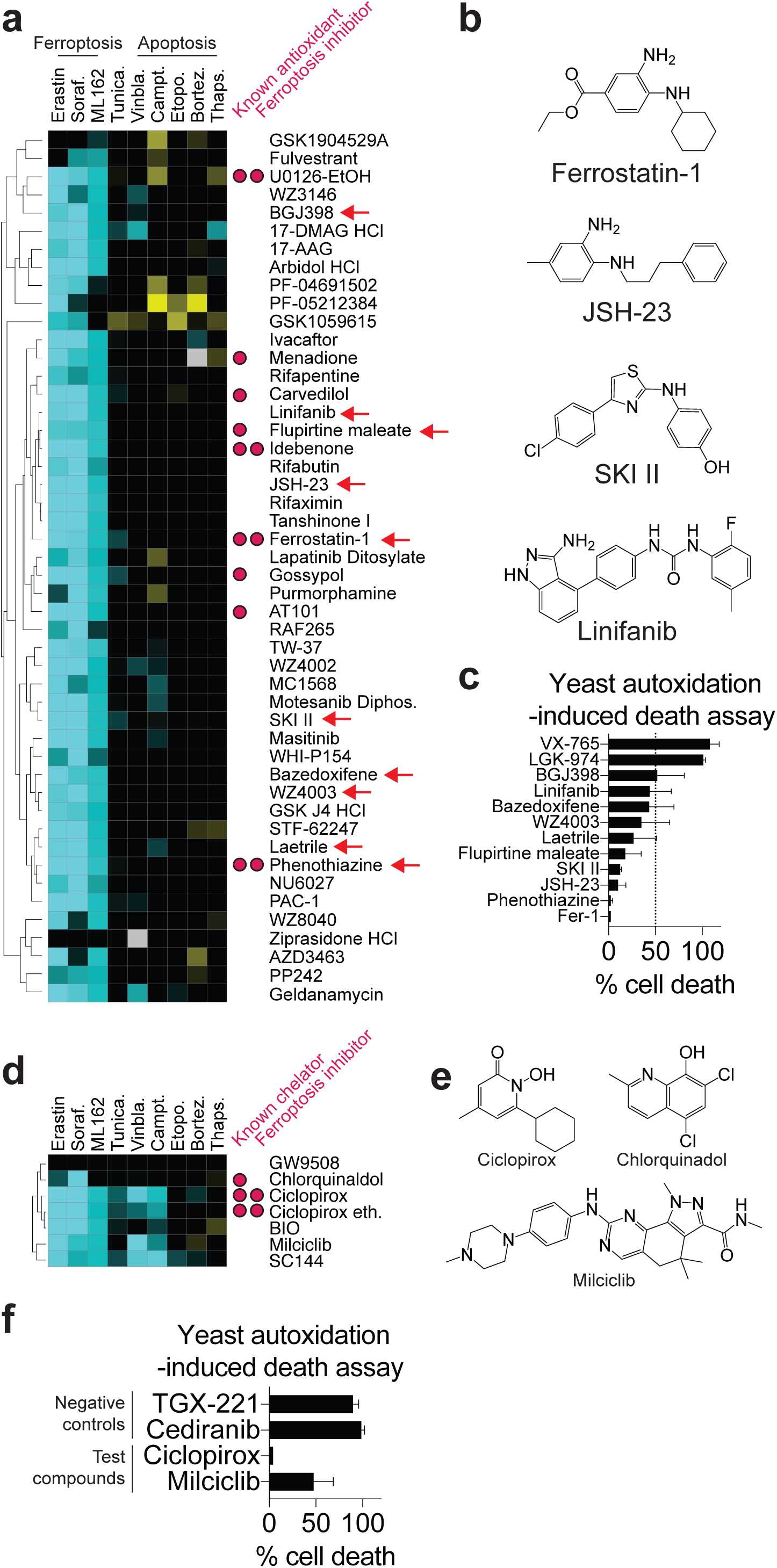
Analysis of ferroptosis-modulating compound clusters. **a**, A cluster of compounds including known ferroptosis inhibitors and known antioxidants without previous associations to ferroptosis (green bar in **Fig. 1c**). Arrows indicate compounds tested in **c**. **b**, Structures of selected compounds within the cluster in **a**. **c**, Cell death in a yeast (*S. cerevisiae*)-based assay for lipid autoxidation-induced cell death. All compounds were tested at 10 µM. VX-765 and LGK-974 were randomly selected negative controls. **d**, A cluster of compounds including the known iron-chelating ferroptosis inhibitors. **e**, Structure of known and candidate iron chelators. **f**, Cell death in the yeast lipid autoxidation-induced cell death assay. All compounds were tested at 10 µM. Results in **c** and **f** represent mean ± SD from three independent experiments.

To test whether compounds in this cluster exhibited previously undescribed antioxidant effects we examined the ability of eight randomly-selected candidates (BGJ398, linifanib, bazedoxifene, WZ4003, laetrile, flurpitine maleate, SKI II, JSH-23) to suppress cell death in a yeast-based assay for polyunsaturated fatty acid autoxidation-induced cell death^40^, independent of modulation by glutathione or GPX4 (**Supplementary Fig. 3b**). All eight candidate compounds, along with two known positive control RTAs (Fer-1, phenothiazine), inhibited yeast autoxidation-induced cell death by at least 50%, while two randomly-selected library compounds (VX-765, LGK-974) did not (**Fig. 2c**). Thus, intrinsic antioxidant activity is likely to explain the ability of these compounds to inhibit ferroptosis.

In addition to putative antioxidants, we observed a distinct cluster of pan-ferroptosis suppressors centered on the iron chelator ciclopirox. This cluster also included chlorquinaldol, which contains the known metal chelating scaffold 8-hydroxyquinolone^41^, suggesting that this may represent a cluster of iron chelating compounds (**Fig. 2d,e**). The cyclin dependent kinase inhibitor milciclib was a novel predicted metal chelator and we confirmed that this compound blocked cell death in the yeast autoxidation assay, consistent with the possibility that it acted as an iron chelator (**Fig. 2f**). These results suggested that cryptic radical trapping and iron chelation effectors might be more common amongst bioactive compounds than previously anticipated.

### Cell-free profiling of free radical scavenging and iron chelation

The above results suggested it would be useful to step back and examine more systematically the extent of unanticipated chemical reactivities in our bioactive compound library. We hypothesized that additional RTAs or iron chelating compounds could exist that were not found in the above cluster, possibly because of multiple actions in cells or lower potency. Thus, to systematically examine the extent of intrinsic free radical scavenging or iron chelating activity amongst bioactive compounds we profiled the 1,833-member compound library for both cell-free antioxidant and iron chelator activity using the 2,2-diphenyl-1-picrylhydrazyl (DPPH) free radical scavenging assay^2^, and the ferrozine Fe^2+^ binding assay^42^, respectively. 5.2% (95/1818) and 6.6% (110/1671) of all assayable bioactive compounds reduced signal by ≥ 50% in the DPPH and ferrozine assays, respectively, with eight compounds active in both assays (**Fig. 3a**).

**Figure 3.**
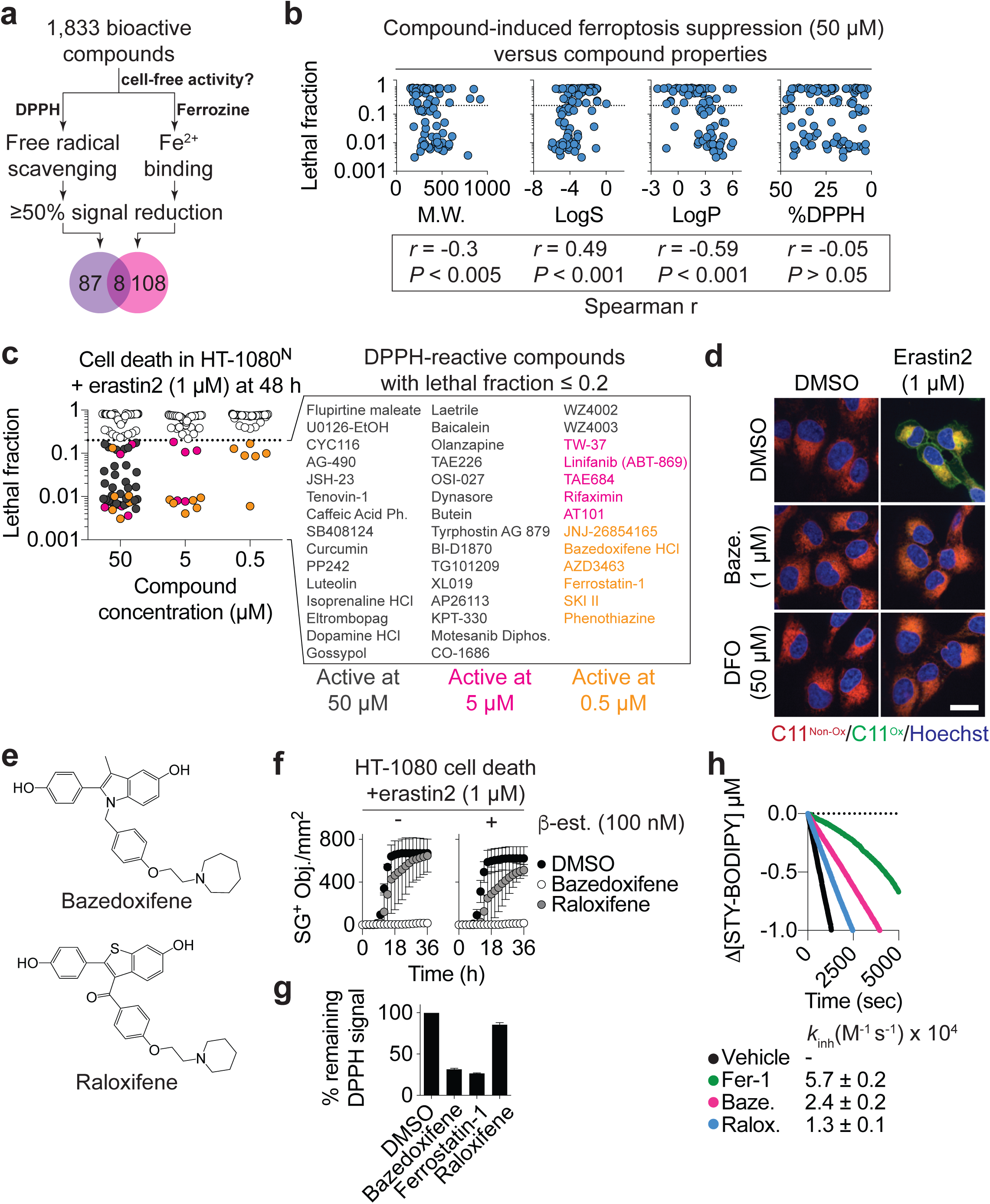
Widespread bioactive compound antioxidant and iron chelating activity. **a**, 1,833 bioactive compounds profiled for cell-free radical scavenging and Fe^2+^-binding activity. **b**, HT-1080^N^ cells were treated with erastin2 (1 µM) plus each of the 95 DPPH-positive compounds (50 µM) and the lethal fraction was determined at 48 h. For each compound, lethal fraction scores are plotted against compound molecular weight (M.W.), predicted hydrophilicity (LogS), predicted lipophilicity (LogP) and the original %DPPH reduction values from the cell-free assay. **c**, The ability of 100 candidate compounds (95 from **b** plus five additional candidates) to suppress erastin2-induced cell death. 46 compounds that suppressed cell death by > 80% are listed, color coded by potency. Ph.: phenethyl ester; Diphos.: diphosphate. **d**, Lipid ROS detected using C11 BODIPY 581/591 in HT-1080 cells. Baze.: bazedoxifene; DFO: deferoxamine; Non-ox: non-oxidized; Ox: oxidized. Scale bar = 20 µm. Imaging was performed twice and representative images from one experiment are shown. **e**, Structures of bazedoxifene and raloxifene. **f,** Effect of raloxifene and bazedoxifene (both 1 µM) on erastin2-induced cell death in HT-1080^N^ cells ± β-estradiol (100 nM). Cell death was quantified as the number of SYTOX Green positive (SG^+^) objects (i.e. dead cells) over time. Data represent mean ± SD from two independent experiments. **g**, Free radical scavenging in the DPPH assay. Data are plotted as the percent remaining signal relative to the negative control (DMSO), which was normalized to 100%. Data are mean ± SD from at least three independent experiments. **h**, Kinetic competition arising in the inhibited co-autoxidation of egg-phosphatidylcholine (1 mM) and STY-BODIPY (10 µM) initiated with 0.2 mM di-tert-undecyl hyponitrite (DTUN). Co-autoxidation (black trace, negative control), inhibited by 2 µM of ferrostatin-1 (green, positive control), bazedoxifene (pink) or raloxifene (blue). Average inhibition rate constants (*k*_inh_) determined assuming a reaction stoichiometry (*n*) of unity.

Subsequently in this work we focused on the 95 compounds that scored positive in the DPPH assay; detailed investigation of compounds positive in the ferrozine assay will be reported elsewhere. These 95 compounds, along with five additional compounds that just missed the 50% threshold but that were included in the subsequent analysis, provided a test set with which to search for chemical properties of free-radical scavenging compounds that might correlate with effective ferroptosis suppression in cells. Accordingly, we tested the ability of all 100 compounds to suppress erastin2-induced ferroptosis in HT-1080^N^ cells when tested at a high concentration (50 µM) and computed chemical properties of interest from each structure. Ferroptosis suppression was significantly correlated with higher compound molecular weight, lower predicted water solubility (LogS) and higher predicted lipophilicity (LogP). The observed relationship across tested compounds between ferroptosis inhibitory potency and LogP is consistent with our previous examination of the isolated Fer-1 structure-activity relationship, which showed the same trend^2^. Notably, the ability of these 100 compounds to suppress erastin2-induced ferroptosis did not correlate with activity in the cell-free DPPH radical scavenging activity (**Fig. 3b**). This is however consistent with the recent finding that there is no correlation between the DPPH radical scavenging activity of 28 putative antioxidants and their ability to suppress RSL3-induced ferroptosis^43^. Thus, while DPPH does provide a useful first pass to detect candidate RTAs, it is insufficient to predict the potency of these molecules against ferroptosis.

Many of these 100 DPPH-positive compounds were also found in the antioxidant cluster, and we next investigated the relative ferroptosis-inhibitory potency of each molecule. 43, 11, and 6 compounds suppressed erastin2-induced ferroptosis by at least 80% (i.e. lethal fraction ≤ 0.2) at 50 µM, 5 µM and 0.5 µM, respectively (**Fig. 3c**). The six most potent compounds, ferrostatin-1, SKI II, phenothiazine, the HDM2/MDM2 inhibitor JNJ-26854165 (serdemetan), the anaplastic lymphoma kinase inhibitor AZD3463, and the third-generation selective estrogen receptor (ER) modulator (SERM) bazedoxifene, all suppressed both erastin2 and ML162-induced ferroptosis with EC_50_ values below 110 nM in confirmatory dose-response experiments, and no effect on direct H_2_O_2_-induced cell death (**Table 1**). Of the most potent ferroptosis inhibitors, JNJ-26854165 was the only compound not found in the original antioxidant cluster (see **Fig. 2a**). It is possible that JNJ-26854165 clustered elsewhere in the original modulatory profile due to additional effects on cell death (e.g. p53 stabilization) unrelated to radical scavenging. This highlights the value of integrating both cell-based and cell-free assays into this analysis.

**Table 1.**
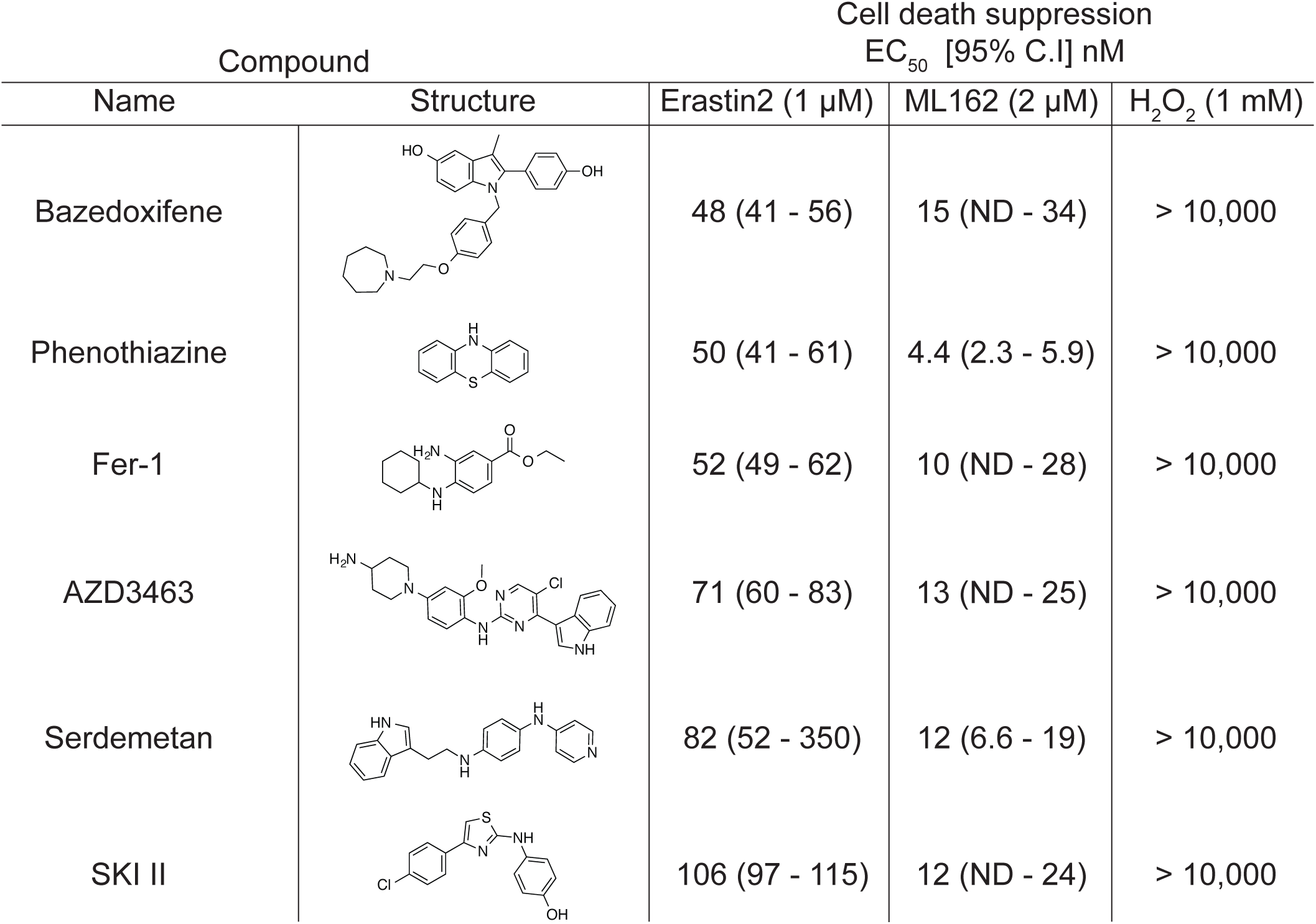
Dose-response testing of ferroptosis inhibitors identified as DPPH free radical scavengers. Name, structure and inhibitor potency in cell-based assays. HT-1080^N^ cells were treated with lethal compounds (erastin2, ML162 or H_2_O_2_) at a fixed concentration and cotreated with a two-point, two-fold concentration-response series of each inhibitor, starting at 10 µM. Lethal fraction data derived from STACK analysis at the 24 h timepoint were used to compute EC_50_ values. Data are mean ± 95% confidence interval (95% C.I.) from three independent experiments. ND: not defined.

| Compound |  | Cell death suppression<br>EC<sub>50</sub> [95% C.I.] nM |  |  |
| --- | --- | --- | --- | --- |
| Name | Structure | Erastin2 (1 µM) | ML162 (2 µM) | H<sub>2</sub>O<sub>2</sub> (1 mM) |
| Bazedoxifene |  | 48 (41 - 56) | 15 (ND - 34) | > 10,000 |
| Phenothiazine |  | 50 (41 - 61) | 4.4 (2.3 - 5.9) | > 10,000 |
| Fer-1 |  | 52 (49 - 62) | 10 (ND - 28) | > 10,000 |
| AZD3463 |  | 71 (60 - 83) | 13 (ND - 25) | > 10,000 |
| Serdemetan |  | 82 (52 - 350) | 12 (6.6 - 19) | > 10,000 |
| SKI II |  | 106 (97 - 115) | 12 (ND - 24) | > 10,000 |

We were intrigued by the ability of bazedoxifene, an FDA-approved drug, to potently suppress ferroptosis. Indeed, we found that bazedoxifene effectively suppressed the accumulation of lipid hydroperoxides in HT-1080 cells treated with erastin2, much like the positive control Fer-1 (**Fig. 3d**). Out of 29 estrogen or progesterone receptor modulators profiled in the compendium, only one other, the structurally-related molecule raloxifene, suppressed ferroptosis to a similar extent as bazedoxifene (**Fig. 3e**, **Supplementary Fig. 3a**). However, in confirmatory studies raloxifene inhibited ferroptosis less potently over time than bazedoxifene when tested at 1 µM, and this did correlate with reduced activity in the DPPH assay (**Fig. 3g**). However, in light of our global findings above concerning the limitations of the DPPH assay, and to provide further insight, we determined the reactivities of bazedoxifene and raloxifene toward phospholipid-derived peroxyl radicals using a recently-developed liposome-based hyponitrite-initiated co-autoxidation assay (**Supplementary Fig. 3c**)^43^. This approach previously revealed an excellent correlation between RTA activity and protection against RSL3-induced ferroptosis^43^. Consistent with this, we observed that bazedoxifene (*k*_inh_ = 2.4×10^4^ M^-1^s^-1^) is roughly twice as reactive as raloxifene (*k*_inh_= 1.3×10^4^ M^-1^s^-1^) toward phospholipid-derived peroxyl radicals when both compounds were tested at 2 µM (**Fig. 3h**). Of note, the protective effect of bazedoxifene was not lost in the presence of the ER antagonist 17β-estradiol, further supporting the notion that it is independent of ER functional modulation (**Fig. 3f**). Together, these results pinpoint numerous small molecules with RTA or iron-binding activity, increase the number of ferroptosis inhibitor compounds, and identify candidate drug repurposing opportunities.

### mTOR inhibition suppresses ferroptosis through an on-target mechanism

We next sought to identify on-target modulators of ferroptosis sensitivity. Intriguingly, within the cell death compendium we observed a cluster of eight ATP-competitive mTOR inhibitors (‘TORKis’) and dual mTOR/PI3K inhibitors that potently suppressed ferroptosis induced by the system x_c_^-^ inhibitors erastin or sorafenib, but not the GPX4 inhibitor ML162 or most pro-apoptotic compounds (**Fig. 4a**). These results suggested that these mTOR inhibitors were not likely acting as antioxidants or iron chelators, which would also be expected to inhibit ML162-induced cell death. Indeed, none of these compounds were identified to possess cell-free RTA or iron scavenging activity. In the original modulatory profile, each compound was tested at 5 µM, a concentration far higher than typically used with mTOR inhibitors. We confirmed that the TORKis INK128 and AZD8055 suppressed erastin2-induced ferroptosis and phosphorylation of the mTOR target 4E binding protein 1 (4EBP1) in HT-1080^N^, U-2 OS^N^ and human embryonic kidney (HEK) 293T^N^ cells when tested at a lower concentration (1 µM) (**Fig. 4b,c**). Unlike mTOR and dual mTOR/PI3K inhibitors, inhibitors of PI3K and AKT had no ability to inhibit ferroptosis (**Supplementary Fig. 4a-d**). Thus, mTOR inhibition was specifically associated with reduced sensitivity to ferroptosis induced by system x_c_^-^ inhibition.

**Figure 4.**
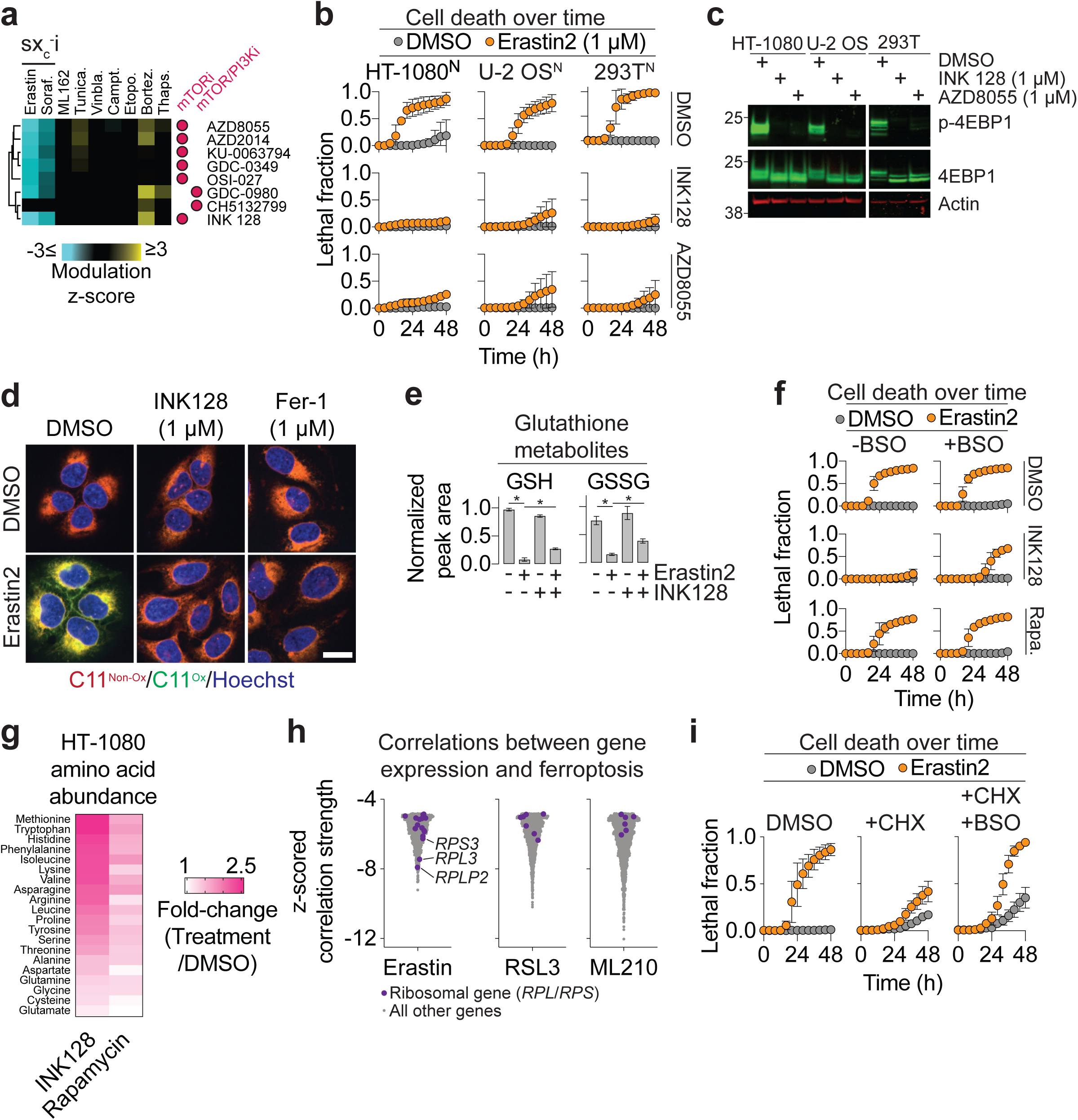
mTOR inhibition suppresses ferroptosis. **a**, A sub-cluster of mTOR and dual mTOR/PI3K inhibitors, from the larger compendium (purple bar in **Fig. 1c**), that block ferroptosis induced by system x_c_^-^ inhibition (sx_c_^-^i). **b**, Cell death over time measured using STACK in three cell lines. INK128 and AZD8055 were used at 1 µM. **c**, Immunoblotting for phosphorylation and total levels of the canonical mTOR target, 4EBP1. **d**, Lipid ROS detected using C11 BODIPY 581/591. U-2 OS cells were treated for 22 h then lipid ROS were imaged using C11 BODIPY 581/591. Erastin2 was used at 2 µM. Fer-1: ferrostatin-1. Scale bar = 20 µm. Imaging was performed three times and representative images from one experiment are shown. **e**, Relative concentration of reduced (GSH) and oxidized (GSSG) glutathione in HT-1080 cells detected using liquid chromatography coupled to mass spectrometry (LC-MS). Erastin was used at 10 µM and INK128 at 5 µM. *P < 0.05, one-way ANOVA with Bonferroni post-tests (n = 4). **f**, Cell death over time in U-2 OS^N^ cells. Compound concentrations were: erastin2 (2 µM), buthionine sulfoximine (BSO, 1 mM), INK128 (1 µM), rapamycin (Rapa., 100 nM). **g**, Fold-change amino acids levels in HT-1080 cells determined using LC-MS. INK128 was used at 5 µM and rapamycin at 100 nM. **h**, Correlations between gene expression and sensitivity to ferroptosis-inducing agents in the CTRP dataset. Genes encoding ribosomal subunits are colored purple. **i**, Cell death over time in U-2 OS^N^ cells treated ± erastin2 (2 µM) ± cycloheximide (CHX, 10 µg/L) ± BSO (1 mM). Data in **b**, **f** and **i** are mean ± SD from three independent experiments.

Mechanistically, TORKi treatment blocked lipid ROS accumulation in erastin2-treated cells and increased the levels of intracellular reduced (GSH) and oxidized (GSSG) glutathione (**Fig. 4d,e**). The ability of TORKi treatment to inhibit ferroptosis was blunted when cells were cotreated with the de novo GSH biosynthesis inhibitor buthionine sulfoximine (BSO)^44^ (**Fig. 4f**). This suggested that inhibition of mTOR signaling promoted survival through effects on glutathione levels. This was reminiscent of the ability of wild-type p53 stabilization to increase intracellular glutathione and inhibit ferroptosis^45^. However, TORKis delayed erastin2-induced ferroptosis in p53 null H1299^N^ non-small cell lung cancer cells, indicating that p53 stabilization alone could not explain this protective effect (**Supplementary Fig. 4e**). TORKi treatment did not increase system x_c_^-^ activity and could also protect from ferroptosis induced by growth in medium lacking cystine (**Supplementary Fig. 4f,g**). Thus, TORKi treatment did not inhibit ferroptosis by enhancing system x_c_^-^-mediated cystine uptake.

TORKis prevented ferroptosis more effectively than rapamycin and related allosteric mTOR inhibitors (‘rapalogs’) (**Supplementary Fig. 4a,g**). Protein synthesis, autophagy and proteasome function are all more strongly impacted by TORKis than rapalogs^46^, and we hypothesized that these processes could conceivably enhance glutathione levels and thereby inhibit ferroptosis by increasing the availability of precursor amino acids. Consistent with this, INK128 treatment increased the relative abundance of most amino acids detected by liquid chromatography and mass spectrometry, while rapamycin had more modest effects (**Fig. 4g**). We previously showed that genetic silencing of *RPL8*, encoding a ribosomal large subunit protein, suppressed ferroptosis induced by erastin but not RSL3 (Ref.^2^). Consistent with this result, across hundreds of cancer cell lines in the Cancer Therapeutics Response Portal (CTRP) dataset^47^, we found more (and stronger) correlations between ribosomal subunit gene expression and sensitivity to erastin (e.g. for *RPS3*, *RPL3*, *RPLP2*), than for sensitivity to the GPX4 inhibitors RSL3 and ML210 (**Fig. 4h**). We hypothesized that mTOR-driven protein synthesis promoted ferroptosis in response to system x_c_^-^ inhibition by consuming precursor amino acids that could otherwise be used for GSH synthesis. This model would account for the observation that direct protein synthesis inhibition (e.g. using cycloheximide) can suppress ferroptosis induced by erastin but not GPX4 inhibitors^48^. Indeed, the ability of CHX to suppress erastin2-induced ferroptosis in U-2 OS^N^ cells was largely reverted by cotreatment with BSO (**Fig. 4i**). These results are consistent with the model that reduced protein synthesis contributes to protects against ferroptosis in response to system x_c_^-^ inhibition by increasing the pool of precursors available for GSH synthesis. Other mTOR-regulated processes likely also contribute to the protective effect of mTOR inhibition. For example, co-treatment with the lysosomal deacidifier chloroquine, which inhibits autophagy, partially attenuated the protective effect of TORKi treatment on ferroptosis (**Supplementary Fig. 4h**). Collectively, these results show that an attenuation of mTOR activity helps the cell maintain intracellular GSH levels, which is essential to prevent ferroptosis following system x_c_^-^ inhibition.

### mTOR-independent regulation of ferroptosis sensitivity by amino acid uptake

Our cell death modulatory profile pointed towards synthetic mTOR inhibitors as potent suppressors of ferroptosis induced by system x_c_^-^ inhibition. We sought to extend this result beyond small molecule inhibitors, showing that manipulation of amino acids levels could directly impact ferroptosis sensitivity. mTOR pathway activity is sensitive to amino acid levels, especially leucine (Leu) and arginine (Arg)^49^. We therefore tested whether depriving cells individually of either Leu or Arg was sufficient to suppress erastin2-induced ferroptosis. Medium lacking Leu had no effect on ferroptosis (**Fig. 5a**). By contrast, medium lacking Arg potently suppressed ferroptosis and blocked lipid ROS accumulation in U-2 OS cells (**Fig. 5a-c**). This protective effect was seen in seven cancer cell lines from diverse lineages and could be reverted by re-supplementing cells with L-Arg or the Arg metabolic precursor citrulline, but not D-Arg (**Supplementary Fig. 5a-c**). Cells deprived of Arg could conserve a small amount of glutathione in response to erastin2 treatment compared to cells grown in complete medium (**Supplementary Fig. 5d**). Functionally, the ability of Arg deprivation to inhibit ferroptosis was reverted by co-incubation with BSO, suggesting that the protective effect of Arg deprivation requires de novo GSH synthesis (**Fig. 5d**).

**Figure 5.**
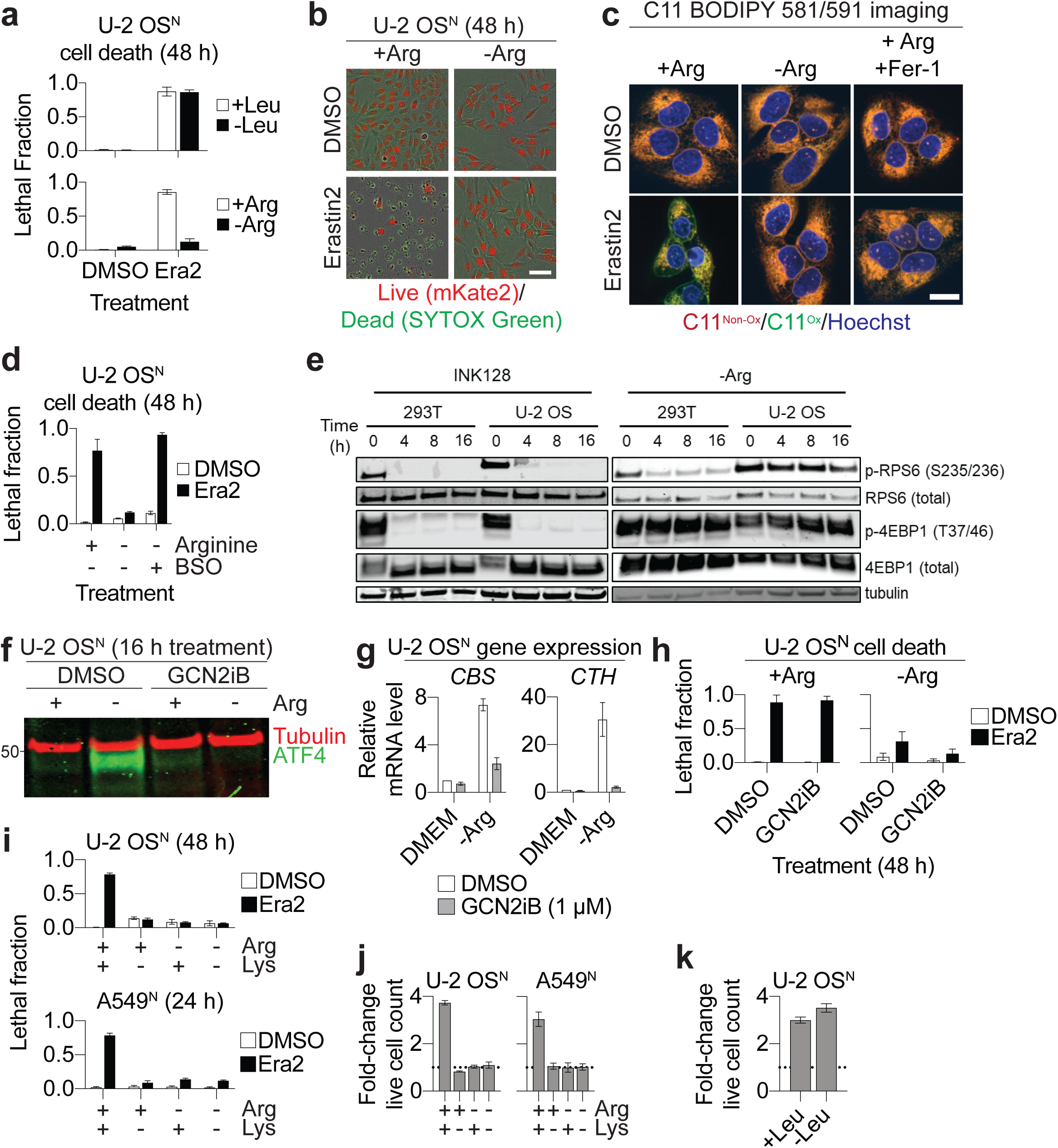
Arginine deprivation inhibits ferroptosis. **a**, Cell death in medium containing dialyzed serum and formulated to lack arginine (Arg) or leucine (Leu). Erastin2 (Era2) was used at 2 µM. **b**, Images of cells at 48 h. Scale bar = 25 µm. **c**, Lipid ROS detected in C11 BODIPY 581/591. U-2 OS cells were treated as indicated for 24 h prior to imaging. Erastin2 was used at 2 µM, ferrostatin-1 (Fer-1) at 1 µM. Non-ox: non-oxidized; Ox: oxidized. Scale bar = 20 µm. Imaging was performed twice and representative images from one experiment are shown. **d**, Cell death at 48 h. Compound concentrations were: erastin2 (2 µM), BSO (1 mM). **e**, Immunoblotting for the mTOR readouts phospho-RPS6 (p-RPS6) and phospho-4EBP1/2 (p-4EBP1/2). INK128 was used at 1 µM. **f**, Model of the transsulfuration pathway. PPG: propargylglycine. **f**, Western blot of ATF4 protein levels ± Arg-containing medium ± the GCN2 inhibitor GCN2iB (1 µM). **g**, Expression of the transsulfuration pathway genes *CBS* and *CTH* under the same conditions as in **f**. **h**, Cell death in U-2 OS^N^ cells at 48 h ± Arg-containing medium ± erastin2 (Era2, 2 µM) ± the GCN2 inhibitor GCN2iB (1 µM). **i**, Cell death ± erastin2 (Era2, 2 µM) in medium lacking either or both Arg and lysine (Lys). **j,i**, Cell proliferation as determined from the ratio of live cell (mKate2^+^) counts at 48 h versus 0 h, in different cell lines and medias ± Arg, Lys or Leu. Data in **a**, **d** and **g**-**k** are mean ± SD from three independent experiments.

These data were consistent with the expectation that Arg deprivation blocked ferroptosis by inhibiting mTOR activity, as seen with TORKis. Indeed, Arg deprivation reduced the phosphorylation of the canonical mTOR target RPS6 in 293T cells, as expected^50^. However, unlike treatment with INK128, Arg deprivation did not inhibit the phosphorylation of RPS6 or 4EBP1 in U-2 OS cells (**Fig. 5e**). Consistent with these observations, bulk protein synthesis was reduced by INK128 treatment but not by Arg deprivation over 16 h, as determined using a puromycylation assay (**Supplementary Fig. 5e**). Thus, Arg deprivation suppressed ferroptosis without inhibiting mTOR activity.

We considered the possibility that Arg deprivation inhibited erastin2-induced ferroptosis by activating the amino acid sensing GCN2/ATF4 pathway, and upregulating transsulfuration metabolism, an alternative route to cysteine synthesis^51–53^. Arg deprivation increased ATF4 protein expression and expression the transsulfuration pathway genes *CBS* and *CTH* (**Fig. 5f,g**). These effects were largely abrogated by co-treatment with the specific GCN2 inhibitor GCN2iB^51^ (**Fig. 5f,g**). However, GCN2iB cotreatment did not prevent Arg deprivation from inhibiting ferroptosis (**Fig. 5h**). Thus, increased GCN2/ATF4 and/or transsulfuration pathway activity could not explain the ability of Arg deprivation to suppress ferroptosis. Note that GCN2iB was unlikely to act as an RTA, as it had no ability to prevent erastin2-induced ferroptosis in regular medium. One possibility was that Arg metabolism specifically promoted ferroptosis through the production of a unique metabolite; however, this seemed unlikely as both U-2 OS^N^ and A549^N^ cells were also protected against ferroptosis by lysine deprivation (**Fig. 5i**). Rather, we noted that both Arg and Lys deprivation cause growth arrest of the cells (**Fig. 5j**). By contrast, Leu deprivation, which did not inhibit ferroptosis, did not arrest cell proliferation (**Fig. 5a,k**). Thus, growth arrest may be sufficient to protect from ferroptosis independent of mTOR inhibition or GCN2/ATF4 pathway activation.

## Discussion

Here, we generated a compendium of time-resolved population cell death profiles and identified dozens of modulators of ferroptosis and other cell death pathways. We also uncover unexpected compound-compound interactions, many of which are best explained by compound off-target chemical reactivities. Examination of this compendium revealed many compounds that inhibit ferroptosis by acting as RTAs or iron chelators. A number of putative lipoxygenase inhibitors were recently shown to inhibit ferroptosis by acting as RTAs^52^. Our study indicates that such effects are more widespread than previously appreciated, and highlight FDA-approved drugs, such as bazedoxifene, as potential drug repurposing candidates to inhibit pathological ferroptotic cell death in the brain, kidney and potentially other tissues, a critical unmet medical need^10^. Bazedoxifene is an interesting, and rare, example of a potent phenolic RTA inhibitor of ferroptosis^43^. The greater RTA potency of bazedoxifene relative to the related molecule raloxifene could be explained by the weaker O-H bond in the 5-hydroxyindole moiety in bazedoxifene compared to the 6-hydroxybenzothiophene moiety in raloxifene, which also contains an electron-withdrawing carbonyl substituent at the C3 position (see discussion in^53^). Further examination of this compendium may reveal additional useful small molecule modulators of cell death. More broadly, this modulatory profiling approach, which incorporates direct measurement of live and dead cells, may be useful to identify new functional associations, make novel compound mechanism of action predictions, and further explore the basic nature of compound-compound interactions over time in mammalian cells^32, 54^.

Balancing amino acid supply with demand is crucial to maintain cell viability. The amino acid cysteine is used for protein synthesis and the synthesis of GSH. ATP-competitive mTOR inhibitors attenuated GSH depletion and delayed the onset of ferroptosis under conditions of cystine limitation (e.g. due to system x_c_^-^ inhibition). We propose that mTOR inhibition, by reducing protein synthesis and increasing autophagy, allows for intracellular Cys levels to be increased and used preferentially for GSH synthesis. This effect would prolong cell survival when extracellular cystine is no longer being imported. This model is also consistent with evidence that protein synthesis normally consumes the majority of intracellular cysteine^55^, and with evidence that oxidative cell death can be rescued by shunting Cys reserves into GSH synthesis^56^. Our results also strengthen the notion that mTOR signaling can have a paradoxical lethal effect, by promoting biosynthetic processes that consume precious resources necessary for survival^57, 58^. Inhibition of the mTOR pathway could be explored as a means to delay ferroptosis in neurodegeneration or other pathological contexts^10^.

Cystine deprivation is a potent trigger for ferroptosis. Intriguingly, induction of this process in response to cysteine deprivation requires the presence of other amino acids, including Arg and Lys. This requirement does not appear to be linked to mTOR pathway activation or suppression of GCN2/ATF4 pathway activity *per se*. Rather, our results suggest that additional mechanisms may exist that can sense the levels of certain amino acids and trigger proliferative arrest when these levels drop below a certain threshold. Here, proliferative arrest correlates with protection from ferroptosis, possibly by slowing metabolism to limit the consumption of GSH and precursor amino acids. These results indicate that a careful balance of amino acids is required for the execution of ferroptosis, and conditions where Arg, Lys or potentially other amino acids are low (e.g. in the tumor microenvironment)^59^ may create resistance to the induction of ferroptosis by cysteine deprivation. This could explain why extracellular cyst(e)ine-degrading enzyme treatment slow but do not shrink tumors in vivo^60, 61^.

## Acknowledgments

We thank J. Cao, A. Tarangelo, Z. Inde and E. Kolebrander Ho for experimental assistance; M. Bassik for H23^Cas9^ cells, and M. Cyert for the *coq3*Δ yeast strain. Certain constructs were obtained from Addgene. This work was supported by the NIH (4R00CA166517-03, 1R01GM122923), NSERC (RGPIN-06741-2016) and a Damon Runyon-Rachleff Innovation award to S.J.D.

## Author contributions

M.C., C.P., G.C.F., L.M., D.A.P. and S.J.D. designed experiments. M.C., C.P., G.F., A.K., M.M., and L.M. performed experiments and analyses. S.J.D. wrote the paper.

## Methods

### Chemicals and reagents

An 1,833-member bioactive compound library (Cat No. L1700), and an independent 86-member PI3K signaling inhibitor library (Cat No. L2800) comprising mTOR, PI3K and AKT pathway inhibitors, were obtained from Selleck Chemicals (Houston, TX) and stored at −80°C. Erastin2 (compound 35MEW28 in Ref.^3^) and ML162 were synthesized by Acme Bioscience (Palo Alto, CA, USA). Erastin was the kind gift of B. Stockwell (Columbia). Chloroquine (Cat No. C6628), ferrostatin-1 (Cat No. SML0583), thapsigargin (Cat No. T9033), tunicamycin (Cat No. T7765), cycloheximide (Cat No. C7698), L-arginine (Cat No. A5131), D-arginine (Cat No. A2646), L-citrulline (Cat No. C7629) and DL-propargylglycine (Cat No. P7888) were obtained from Sigma-Aldrich Corporation. Bortezomib (Cat No. NC0587961), buthionine sulfoximine (Cat No. AC23552-0010), were obtained from Thermo-Fisher Scientific. INK 128 (Cat No. S2811), AZD8055 (Cat No. S2661), vinblastine (Cat No. S1248), camptothecin (Cat No. S1288), sorafenib (Cat No. S7397), bazedoxifene (Cat No. S2128), raloxifene (Cat No. S1227) and JTC-801 (Cat No. S2722) were obtained from Selleck Chemicals. Rapamycin (Cat No. BP2963-1) and etoposide (Cat No. 12-261-00) were obtained from Fisher Scientific. GCN2iB was from MedChemExpress (Cat No. HY-112654). Buthionine sulfoximine and DL-propargylglycine were dissolved directly into cell media. All other drugs were prepared as stock solutions in DMSO. Stock solutions were stored at −20°C.

### Cell culture

HT-1080 (CCL-121), U-2 OS (HTB-96), HEK293T (CRL-3216, hereafter 293T), NCI-H1299 (CRL-5803, hereafter H1299), A549 (CCL-185), T98G (CRL-1697), Caki-1 (HTB-46) and A375 (CRL-1619) were obtained from ATCC (Manassas, VA, USA). H23 cells stably-expressing Cas9 (H23^Cas9^) were the kind gift of Michael Bassik (Stanford). The polyclonal nuclear mKate2-expressing (denoted by superscript ‘N’) cell lines HT-1080^N^, U-2 OS^N^, 293T^N^ and H1299^N^ were described previously^20, 45^. Polyclonal populations of Caki-1^N^, A375^N^ and H23^Cas9,N^ cells were generated from the respective parental cells via transduction with the NucLight Red lentivirus, which directs the expression of nuclear-localized mKate2 (Cat No. 4476, Essen BioSciences/Sartorius, Ann Arbor, Michigan, USA). Polyclonal mKate2-expressing populations were selected using puromycin (Life Technologies, Cat No. A11138-03, 1.5 μg/mL, for 48-72 h). HT-1080 cells were cultured in DMEM Hi-glucose media (Corning Life Science, Cat. No. MT-10-013-CV) supplemented with 1% non-essential amino acids (NEAAs, ThermoFisher Scientific, Cat. No. 11140050). A549, 293T, and T98G cells were cultured in DMEM Hi-glucose medium without supplemental NEAAs. U-2 OS, Caki-1 and A375 cells were cultured in McCoy’s 5A medium (Corning Life Science, Cat. No. MT-10-050-CV). H23^Cas9^ cells were cultured in RPMI 1640 with L-glutamine medium (Fischer Scientific Cat No. SH30027FS). All media was supplemented with 10% FBS and 0.5 mg/mL penicillin-streptomycin (Life Technologies, Cat No. 1037-016) unless otherwise indicated. Cell lines were grown at 37°C with 5% CO_2_ in humidified tissue culture incubators (Thermo Scientific, Waltham, MA, USA).

### Bioactive compound library profiling in cells

Large-scale bioactive compound profiling was performed as described^20^. HT-1080^N^ cells were grown in T-175 flasks (Cat. No. 431080, Corning Life Sciences) were trypsinized and counted using a Cellometer Auto T4 cell counter (Nexcelom, Lawrence, MA, USA). 40 μL of cell solution was added manually to each well of a 384 well clear bottom tissue culture plate (Cat No. 3712, Corning) at a final density of 1,500 cells per well. The plate was spun briefly (500 rpm, 2 sec) to settle the cells evenly at the bottom of the wells. The next day, the medium was removed and replaced with 36 µL of media containing the dead cell probe SYTOX Green (20 nM, final concentration; Cat No. S7020, Life Technologies) and DMSO or one of ten lethal compounds (final concentration), thapsigargin (12.5 nM), tunicamycin (10 µg/mL), camptothecin (5 µM), etoposide (100 µM), bortezomib (50 nM), vinblastine (0.1 µg/mL), erastin (10 µM), sorafenib (10 µM) and ML162 (5 µM). 4 µL of medium containing one of 1,833 different bioactive library compounds was then added (final concentration 5 µM) using a Versette robotic liquid handler (ThermoFisher Scientific) and plates were immediately transferred to an IncuCyte Zoom dual color live content imaging system (Model 4459, Essen BioScience/Sartorius, Ann Arbor, MI, USA) housed within a Thermo tissue culture incubator (37°C, 5% CO_2_).

Images were acquired using a 10x objective lens in phase contrast, green fluorescence (ex: 460 ± 20, em: 524 ± 20, acquisition time: 400 ms) and red fluorescence (ex: 585 ± 20, em: 665 ± 40, acquisition time: 800 ms) channels. For each well, images (1392 × 1040 pixels at 1.22 μm/pixel) were acquired every 2 h for a variable period of time reflecting the different kinetics of cell death induced by each lethal query compound (time): DMSO control (120 h), thapsigargin (112 h), tunicamycin (96 h), camptothecin (96 h), etoposide (102 h), bortezomib (96 h), vinblastine (96 h), erastin (46 h), sorafenib (46 h) and ML162 (24 h). Automated object detection was performed in parallel to data acquisition using the Zoom software package (V2016A/B) using a routine with the following settings (in parentheses) to count mKate2^+^ objects (Parameter adaption, threshold adjustment: 1; Edge split on; Edge sensitivity 50; Filter area min 20 μm^2^, maximum 800 μm^2^; Eccentricity max 1.0) and SG^+^ objects (Parameter adaption, threshold adjustment: 10; Edge split on; Filter area min 5 μm^2^, maximum 800 μm^2^; Eccentricity max 0.9).

### Compendium data analysis and visualization

Cell death within each population was analyzed using the scalable time-lapse analysis of cell death kinetics (STACK) approach using counts of live (mKate2^+^) and dead (SYTOX Green, SG^+^) cells to compute lethal fraction scores at each timepoint, as described^20^. At this stage, several data quality filters were applied. First, results obtained for ten autofluorescent compounds were removed from all subsequent analyses: nintedanib (BIBF 1120), sunitinib malate, enzastaurin (LY317615), PHA-665752, SB216763, SU11274, idarubicin HCl, TSU-68 (SU6668, orantinib), quinacrine 2HCl and Ro 31-8220 mesylate. Second, we removed from the analysis 16 bioactive library compounds from the bortezomib profile, and 76 bioactive library compounds from the vinblastine profile, as these were determined to have fewer than 50 live mKate2^+^ cells/well at t = 0, which we set arbitrarily as a cut-off to limit the potential impact of cell density effects to compound sensitivity.

For all populations, lethal fraction scores over time were summarized as a single value by computing the area under the curve (AUC) value of the lethal fraction scores over time, using the default settings in Prism 7 (GraphPad Software, San Diego, CA). AUC values vary as a function of time (i.e. for a given lethal stimulus, a longer period of incubation will result in higher AUC value). Thus, to enable a comparative analysis of bioactive compound modulatory effects between the lethal queries that were each observed for different lengths of time (e.g. ML162 was rapidly lethal and was only observed for 24 h, while thapsigargin was slowly lethal and was observed for 112 h), the AUC values for each population were normalized to the maximum possible cell death within the observation period, *u*. Thus, the normalized AUC (nAUC) = AUC^Observed^ ^(time: 0➔^*^u^*^)^/AUC^Max^ ^(time: 0➔^*^u^*^)^. For example, AUC^Max^ for ML162 = 24, while AUC^Max^ for thapsigargin = 112. nAUC values were also computed for each bioactive compound alone from the DMSO screen at different time intervals (e.g. 24 h for ML162, 112 h for thapsigargin). nAUC values for lethal query (‘Q’) and bioactive compound modulators (‘M’) alone were then used to compute the expected nAUC (nAUC^Expect^) using the Bliss independence model (Q + M - [Q*M])^23, 24^. The difference between nAUC^Expect^ and the experimentally observed nAUC value (nAUC^Observed^) were then calculated for each compound combination (Difference = nAUC^Expect^ – nAUC^Observed^). To account in an unbiased way for differences in the overall ‘modulatability’ of each lethal query by the bioactive compound library compounds, all Difference values were z-scored separately for each lethal query across all tested bioactive modulator compounds. These z-scores were hierarchically clustered in an unsupervised manner as a heat map. Hierarchical clustering was performed using default settings and heatmaps were generated using Morpheus (https://software.broadinstitute.org/morpheus/).

### Identification of known antioxidants and iron chelators

To identify known antioxidants literature searches were performed using PubMed (https://www.ncbi.nlm.nih.gov/pubmed) and the search terms “[compound]” AND (“antioxidant” OR “ferroptosis”) on 18 Feb 2018.

### Analysis of ERK phosphorylation

For Western blots of phosphorylated and total ERK, 200,000 HT-1080 cells per well were grown overnight in a 6-well tissue culture dish (Cat. No. 3516, Corning). Cells were washed with 1 mL Hanks Balanced Salt Solution (HBSS, Cat. No. 14025-134, Life Technologies) and the media was replaced with serum-free DMEM High Glucose media containing 1% Pen/Strep. After 16 h of serum starvation, HT-1080 cells were stimulated with serum-containing media ± U0126 (5 μM) or trametinib (250 nM) for 30 min, washed with 1 mL HBSS and lysed in 100 μL of 9 M urea containing protease and phosphatase inhibitor cocktail (Cell Signaling Technology). Samples were split into two 30 μg aliquots and separated on a Bolt 4-12% Bis-Tris Plus SDS gel (Cat. No. BG04120BOX, Life Technologies) for 2 h at 80 volts. The gel was transferred to a nitrocellulose membrane using the iBlot2 system (Cat. No. IB21001, Life Technologies). Membranes were probed with rabbit monoclonal antibodies directed against the diphosphorylated (Thr^202^/Tyr^204^) form of ERK1/2 (p42/44 MAPK, Cat. No. 4372, Cell Signaling Technology) and total ERK1/2 (Cat. No. 4695, Cell Signaling Technology), both used at a dilution of 1:1000. Goat anti-rabbit secondary antibodies (IRDye 680LT, Cat. No. 926-68021) were from LI-COR Biosciences (Lincoln, NE, USA). Membranes were imaged using a LI-COR CLx scanner.

### Yeast strains and medium

The *Saccharomyces cerevisiae* strain used in this study was *coq3*Δ (BY4741 *MAT***a** *his3*Δ*0 leu2*Δ*0 met15*Δ*0 ura3*Δ*0 coq3*Δ::*kanMX4*). *coq3*Δ was grown in YPD (1% yeast extract (Cat No. 212750, BD Biosciences), 2% peptone (Cat No. 211677, BD Biosciences), 2% dextrose (Cat No. M-15722, Fisher BioReagents).

### Yeast fatty acid treatment

The day before the experiment, a single yeast *coq3*Δ colony was used to inoculate 5 mL YPD. The culture was grown overnight (30°C, 160 rpm). The next morning, the culture was diluted to OD_600_ = 0.1 in fresh YPD and incubated at 30°C, 160 rpm for ∼4 h to mid-log phase (OD_600_ = 0.2-0.5). At mid-log phase, cells were harvested and washed two times with 2 volumes of sterile water, resuspended to a final OD_600_ = 0.8 in 0.1 M sodium phosphate buffer pH 6.2/2% glucose + 1 µM SYTOX Green, and 50 μL cell suspension was added to the appropriate wells of a 96-well clear black-side, clear, flat-bottom plate (Cat No. 3904, Costar) (final SYTOX Green = 250 nM).α-linolenic acid (Cat No. 90210, Cayman Chemical) and vehicle (ethanol) were diluted to 1 mM in 0.1 M sodium phosphate buffer pH 6.2/2% glucose and 100 μL of the appropriate mixture was added to the appropriate wells (final fatty acid concentration = 500 µM). Candidate antioxidants, iron chelators and DMSO (Cat No. 276855, Sigma-Aldrich) were diluted to 40 µM in 0.1 M sodium phosphate buffer pH 6.2/2% glucose, and 50 μL of the appropriate mixture was added to the appropriate wells (final antioxidant concentration = 10 µM). Assay plates were incubated for 24 h (30°C, 160 rpm) in a Cytation3 cell imaging multimode reader (BioTek Instruments, Winooski, VT, USA). At 24 h, the SYTOX Green fluorescence was measured on the Cytation3 using ex/em settings of 488/523. The background signal (0.1 M sodium phosphate buffer pH 6.2/2% glucose + 250 nM SYTOX Green only) was subtracted from all samples, and the final percent cell death was determined using the 500 µM α-LA + DMSO condition set to 100% cell death.

### 2,2-diphenyl-1-picrylhydrazyl (DPPH) assay

For the DPPH profiling experiment of the 1, 833 compound library, the stable radical 2,2-diphenyl-1-picrylhydrazyl (DPPH) was dissolved in methanol (MeOH) to a concentration of 53.3 μM. 60 μL of DPPH solution was added to 20 µL of diluted library compound (160 µM). The final concentrations of DPPH and test compounds were 40 μM. Samples were incubated in the dark at room temperature for 30 min. After incubation, absorbance was measured at 517 nm using a Cytation3 multimode reader. Each plate had eight wells with DPPH and vehicle (DMSO) only and eight wells with MeOH only for background subtraction. Each plate was blank subtracted using the average MeOH signal from eight wells and normalized to eight control wells containing DPPH + DMSO only. The entire experiment was performed twice on separate days. 13 compounds had average normalized DPPH signals ≥ 150% of the negative controls and were excluded from the analysis (obatoclax mesylate, crystal violet, clofazimine, indirubin, vitamin B12, daunorubicin HCl, epirubicin HCl, enzastaurin, doxorubicin, idarubicin HCl, GW441756, BIO, and pirarubicin). Of the remaining 1,820 compounds, 5.2% (95/1818) exhibited normalized DPPH signals between 0-50% of the negative controls, with < 20% standard deviation between the two replicates, and were considered primary hits. When the DPPH assay was performed for single compound follow-up experiments, DPPH was dissolved in methanol to a final concentration of 40.2 μM. 498 μL of DPPH solution was added to 2 uL of 10 mM compound dissolved in DMSO. The final concentrations of DPPH and test compounds were 40 μM. Samples were briefly vortexed and allowed to incubate in the dark at room temperature for 60 min. 150 μL aliquots of each DPPH:test compound solution were added to three wells of 96-well clear-bottom tissue culture plates (Cat. No. 3904, Corning) and absorbance at 517 nm was recorded using a Cytation3 cell imaging multimode reader (BioTek). Absorbance at 517 nm was averaged across the three technical replicates, blank (methanol only) subtracted, and normalized to average DPPH absorbance. The entire experiment was performed three times.

### Ferrozine iron chelation assay

The 1,833-member bioactive compound library was examined using the ferrozine assay as follows. 4 µL of each library compound (2 mM) were diluted to a final concentration of 53.3 µM in two steps in H_2_O with a robotic liquid handler (ThermoFisher Scientific). Next, 4 µL of iron (II) chloride (100 µM) were added to 60 μL of each diluted compound and mixed. Finally, 16 μL of 3-(2-pyridyl)-5,6-diphenyl-1,2,4-triazine-p,p’-disulfonic acid monosodium salt hydrate (ferrozine)^42^ (125 µM) was added to 64 µL of compound + iron (II) chloride and mixed in 384-well clear-bottom plates (Cat No. 3712, Corning). The final concentrations of the resulting 80 μL reactions for test compounds, iron (II) chloride, and ferrozine were 40 μM, 5 μM and 25 μM respectively. After incubating in the dark at room temperature for 60 min, absorbance readings were taken at 562 nm using a Cytation3 multimode reader. Absorbance at 562 nm was averaged for two or three independent experimental trials conducted on different days for each compound, blank (ferrozine only) subtracted, and normalized to average ferrozine:iron absorbance.

### Antioxidant mini-library and follow-up analysis

Compounds identified in the DPPH library screen were cherry picked from a fresh library aliquot of the bioactive compound library for testing for the ability to suppress erastin2 (1 µM)-induced cell death in HT-1080^N^ cells. 95 compounds that reduced DPPH signal by ≥50% were selected, along with five additional compounds close to the cutoff (2-methoxyestradiol [50.01], TCID [50.6], epinephrine HCl [50.8], milciclib [50.8] and PYR-41 [51.4]. The library was tested against a fixed concentration of erastin2 (1 µM) at three doses: 50 µM, 5 µM, and 0.5 µM over 48 h, scanning in 2 h intervals. LF scores were computed for each time point to generate nAUC scores for each treatment. Data represent the mean of two independent experiments where all compounds were tested.

### Analysis of cell death with 17β-estradiol competition assay

17β-estradiol is an estrogen receptor (ER) antagonist with ∼10-fold greater affinity to ER than bazedoxifene or raloxifene^62^. HT-1080^N^ cells were seeded into a 384 well plate (Corning, Cat. No. 3712) at a final density of 1,500 cells/well. Cells were treated with erastin2 (1 µM) in the presence or absence of bazedoxifene or raloxifene (both 1 µM). In addition, cells were treated ± 17β-estradiol (100 nM). Counts of live and dead cells were acquired every 2 h for 36 h.

### Lipophilicity prediction

Compound lipophilicity was predicted using the ALGOPS algorithm to predict logP and logS values^63^ (http://www.vcclab.org/lab/alogps/).

### Kinetic analysis of radical-trapping antioxidant activity

All chemicals and solvents were purchased from commercial suppliers and used without further purification unless other indicated. STY-BODIPY and DTUN were prepared as reported previously^43, 64^. Egg phosphatidylcholine liposomes were prepared as previously reported^65^. UV−visible spectra and kinetics were measured on a Cary-100 UV−vis spectrophotometer equipped with a temperature controller unit and a thermostatted 6 × 6 multicell holder. To a cuvette of 2.34 mL of 10 mM phosphate buffered saline (150 mM) at pH 7.4 was added 125 μL of 20 mM stock of 100 nm unilamellar egg phosphatidylcholine liposomes in the same buffer, and the cuvette was placed into the thermostatted sample holder of a UV−visible spectrophotometer and equilibrated to 37 °C. An aliquot (12.5 μL) of a 2.0 mM solution of STY-BODIPY in DMSO was added, followed by 10 μL of a 50 mM solution of DTUN in EtOH, and the solution was thoroughly mixed. The absorbance of the sample at 571 nm was monitored for ca. 20 min to ensure that STY-BODIPY consumption was proceeding at a constant rate, after which 10 μL of a 500 μM solution of the test antioxidant was added. The solution was thoroughly mixed and the absorbance readings resumed. The initial rate and inhibited period were then used to calculate *k*_inh_ and *n* as described^43^.

### Analysis of cell death with mTOR inhibitors

HT-1080^N^, U-2 OS^N^, 293T^N^ and H1299^N^ cells were seeded in clear bottom 12-well plates (Corning, Cat No. 3737) at densities of 75,000, 50,000, 100,000, and 50,000 cells per well, respectively, in 1 mL of medium. The next day, the medium was replaced with fresh medium containing SYTOX Green (final concentration = 20 nM) along with either DMSO or erastin2 (1 µM), INK 128 (1 µM) and AZD8055 (1 µM). Cells were then transferred into the IncuCyte Zoom enclosed within a tissue culture incubator and images were acquired using the 10x objective every 4 h for 48 h. Live (mKate2^+^) and dead (SG^+^) objects (i.e. cells) were counted using the IncuCyte ZOOM Live-Cell Analysis System software with a custom analyzer and the lethal fraction scores were computed over time, as described above. In some experiments, U-2 OS^N^ cells were co-treated ± buthionine sulfoximine (BSO, 1 mM), rapamycin (Rapa., 100 nM) or chloroquine (CQ, 25 µM). In one experiment, ferroptosis was induced in U-2 OS^N^ cells by placing them in medium formulated to lack cystine, as described^66^.

### PI3K library mini-screen

The effects of 86 different small molecule inhibitors of PI3K and related pathways were examined in U-2 OS cells (shown in Supplementary Fig. 4). Cells were seeded in 384-well plates at a density of 1,500 cells/well in 40 µL of medium. The next day, the medium was removed and 36 µL of fresh medium containing SYTOX Green (20 nM final concentration) and the PI3K library compounds were added. The PI3K library compounds were added to final concentrations of 100 nM, 250 nM and 10 nM in separate plates. Then, either immediately, or following a 6 h pre-incubation, 4 µL of 10x erastin solution (10 µM final concentration) was added to cells, and cells were immediately transferred to an IncuCyte Zoom dual color live content imaging system housed within a Thermo tissue culture incubator, as described above, and imaged every 2 h for the ensuing 24 h. Images were acquired using a 10x objective lens in phase contrast and green fluorescence (ex: 460 ± 20, em: 524 ± 20, acquisition time: 400 ms). Counts of SYTOX Green positive (SG^+^) dead cells at 24 h in response to erastin treatment alone were used to compute a normalized cell death for each condition, set equal to 1. The effect of each individual PI3K pathway inhibitor was assessed relative to this baseline, with a value of 0 being equal to no dead cells observed at 24 h. The entire experiment was repeated three times on separate days and the results shown are the average of these three experiments, where each dot represents an individual compound.

### Cell lysis and immunoblotting

Except where noted, adherent cells were washed once in ice cold PBS and lysed in 40 μL ice cold RIPA buffer (50 mM Tris HCl pH 7.5, 150 mM NaCl, 0.1% SDS, 0.5% sodium deoxycholate, 1% triton, plus protease/phosphatase inhibitor cocktail). Lysate was removed from the well, added into 1.5 mL Eppendorf tubes, and sonicated with 10 one-second pulses at maximum amplitude with a Fisher Scientific Model 120 Sonic Dismembrator (Thermo Fisher). Lysates were centrifuged at 12,700 rpm speed at 4°C for 15 min, and the supernatant was isolated to exclude debris. Protein concentration was quantified using the Pierce Microplate BCA Assay Kit (Cat No. 23252 Thermo Fisher). Samples were prepared with 10x Bolt Sample Reducing Agent (Cat No. B0009 Life Technologies) and 4x Bolt LDS Sample Buffer (Cat No. B0007 Life Technologies) and run on Bolt 4-12% Bis-Tris Plus gels (Cat No. BG04120BOX, Life Technologies) at 100V for 1 h and 45 min using Bolt MES running buffer (Thermo Fisher, Cat. No B0002) diluted to 1x. Protein was transferred to iBlot nitrocellulose membranes (Cat No. IB23001, Life Technologies) using the iBlot2 system, using the preset protocol P0. Membranes were blocked in Odyssey Blocking Buffer (Li-Cor, Cat. No. 927-40010) for one hour at room temperature and incubated with primary antibodies. Primary antibodies used were goat anti-actin (I-19, sc-1616, dilution: 1:1000) from Santa Cruz, rabbit anti-RPS6 (2217S, 1:5000), rabbit anti-phospho-RPS6 Ser235/236 (4858S, 1:2000), rabbit anti-4EBP1 (9644S, 1:1000), rabbit anti-phospho-4EBP1 Thr37/46 (9459S, 1:1000), rabbit anti-ATF4 (11815S, 1:1000) and rabbit anti-GAPDH (2118S, 1:500) from Cell Signaling Technologies, rabbit anti-LC3B (NB600-1384AF700, 1:1000) from Novus Biologicals, and mouse anti-tubulin (MS581P1, 1:10,000) from Fisher Scientific. Membranes were probed overnight at 4°C or for 1 hour at room temperature with rocking. Then, membranes were washed three times for seven min in Tris-buffered saline (ISC BioExpress, Cat. No. 0788) with 0.1% Tween-20 (TBST). Secondary antibodies used were IRDye 680RD Donkey anti-Mouse IgG (LI-COR Biosciences, Cat. No. 926-68072), IRDye 680RD Donkey anti-Goat (926-68074), and IRDye 800 Donkey anti-Rabbit (926-32213). Samples were probed with secondary antibodies at 1:15,000 in Odyssey Blocking Buffer (LI-COR, Cat No. 927-40100), diluted 1:1 with TBST for 1 hour at room temperature. Membranes were washed three times for seven min in TBST, then imaged on the Odyssey CLx Imaging System (LI-COR Biotechnology, Lincoln, NE, USA).

### C11 581/591 BODIPY imaging

The day before the experiment, 150,000 HT-1080 or 100,000 U-2 OS cells/well were seeded into 6-well plates that had one 22 mm^2^ glass coverslip in each well. The next day, the cells were treated with the appropriate compound(s) in the appropriate medium for 11 h (erastin2 treatment) in HT-1080 cells, or 22 h (INK 128 treatment) or 24 h (arginine deprivation) in U-2 OS cells. After the completion of treatment time, the treatment medium was removed and then the cells were treated with C11 BODIPY 581/591 (Cat No. D3861, Molecular Probes, Eugene, OR; final concentration = 5 μM) and Hoechst (Cat No. H1399, Molecular Probes; final concentration = 1 µg/mL) dissolved in HBSS and incubated at 37°C for 10 min. After 10 min, the C11 BODIPY 581/591/Hoechst mixture was removed and fresh HBSS was applied to the cells. The cover slips were mounted in 25 μL HBSS onto glass microscope slides. Cells were imaged using a Zeiss Axio Observer microscope with a confocal spinning-disk head (Yokogawa, Tokyo, Japan), PlanApoChromat 63×/1.4 NA oil immersion objective, and a Cascade II:512 electron-multiplying (EM) CCD camera (Photometrics, Tucson, AZ). Images were processed in ImageJ 1.48v.

### Glutamate release assay

Adherent cells were washed twice in cystine uptake buffer (137 mM choline chloride, 3mM KCl, 1 mM CaCl_2_, 1 mM MgCl_2_,5 mM D-Glucose, 0.7 mM K_2_HPO_4_, 10 mM HEPES, 300 μM cystine, pH 7.4). Uptake buffer containing DMSO or Erastin2 (1 μM) was added to cells and incubated for 60 minutes at 37°C. Cell media was then collected and added to a 96-well assay plate. For normalization purposes, cells were trypsinized and cell number was quantified using a Cellometer Auto T4 Bright Field Cell Counter. Glutamate release was detected using the Amplex Red Glutamic Acid/Glutamate Oxidase Assay kit (Thermo Fisher, Cat No. A-12221) per the manufacturer’s instructions. 10 μM H_2_O_2_ and 25 μM L-glutamate were included as positive controls. Fluorescence readings were recorded at ex/em 530/590 on a BioTek Synergy Neo2 multimode reader. Background fluorescence from blank uptake media was subtracted and samples were normalized to cell number.

### Determination of glutathione and amino acid abundance by mass spectrometry

HT-1080 cells were seeded in 15 cm^2^ dishes and treated with ± erastin2 (1 µM) ± INK 128 (5 µM) or rapamycin (100 nM) for 10 h. Cells were trypsinized, harvested, and cell number was quantified using a Cellometer Auto T4 Bright Field Cell Counter. Cells pelleted at 1,500 x g for 5 min, and pellets were frozen at −80°C. Four biological replicates were collected and analyzed by Metabolon (Durham, NC) as described previously^67^.

### Cancer therapeutics response portal analysis

We obtained data from the Cancer Therapeutics Response Portal (CTRP) v2.1 at https://ocg.cancer.gov/ctd2-data-project/broad-institute-screening-dependencies-cancer-cell-lines-using-small-molecules-0. The data for statistically significant Pearson correlations between basal gene expression and sensitivity to erastin, RSL3 (denoted 1S,3R-RSL-3 in the dataset) and ML210 from all cancer cell lines available for analysis were extracted from the v21.data.gex_global_analysis.txt table and plotted using Prism 8.2.1.

### Amino acid deprivation and resupplementation

All amino acid deprivation experiments were conducted by treating cells in starvation media, resupplemented with stock solutions of the missing amino acid. DMEM minus arginine was constituted by supplementing DMEM for SILAC (Cat No. 88364 Thermo Fisher), which lacks L-lysine and L-arginine, with a 1000x stock solution of L-lysine*HCl at 130 g/L. L-lysine is found in stock DMEM at 0.146 g/L, but once fully diluted with FBS and P/S, DMEM constitutes only 89% of the final volume. Thus, a 1000x stock is 89% of 146 g/L, or 130 g/L. Similarly, DMEM minus leucine was constituted by supplementing DMEM-LM (Cat No. 30030 Thermo Fisher), which lacks L-leucine and L-methionine, with 1000x stock solution of L-methionine at 26.7 g/L, 89% of the 30 mg/L contained in DMEM. To minimize the contribution of monomeric amino acids contained in normal FBS, all medias were supplemented with 10% dialyzed FBS (dFBS, Cat No. 26400044 Thermo Fisher) and 1% P/S.

### Amino acid deprivation and resupplementation cell death analysis

The effects of amino acid deprivation on cell death were investigated in several ways. Most experiments used U-2 OS^N^ cells. These cells were seeded overnight at either 20,000 cells/well in 24-well plate (BD Falcon, Cat No. 353047), 40,000 cells/well in 12-well plates (Corning, Cat No. 3513), or 100,000 cells/well in 6-well plates (Corning, Cat No. 3516), in McCoy’s 5A medium. The next day, cells were washed once in pre-warmed HBSS, the medium was then replaced with DMEM-base resupplementation medium, as described above, containing different treatment conditions, and incubated in a Thermo tissue culture incubator (37°C, 5% CO_2_) for 48 h. Treatment conditions were medium lacking Arg only, medium lacking leucine only, medium lacking Arg and resupplemented with L-Arg, D-Arg or L-citrulline (all at 356 µM), medium lacking Arg resupplemented with increasing concentrations of L-Arg to enable the calculation of the L-Arg EC_50_ (using four-point logistic regression in Prism 7), and medium lacking Arg and supplemented with buthionine sulfoximine (BSO, 1 mM) or DL-propargylglycine (PPG, 2 mM). Both BSO and PPG were dissolved directly into the medium. In one experiment we examined the effects on erastin2-induced cell death of Arg starvation in HT-1080^N^, T98G^N^, A549^N^, Caki-1^N^, H1299^N^, H23^Cas9,N^, U-2 OS^N^ and A375^N^ cells, all seeded in 24-well plates at 20,000 cells/well. In all experiments, images were collected every 4 h for 48 h treatment, live and dead cells were counted using the IncuCyte Zoom dual color imaging system housed within a Thermo tissue culture incubator, and percent ferroptosis suppression was computed as the ratio of AUC values for cells treated with erastin2 and grown with or without arginine. Note that in the resultant graph, values closer to 100% indicate greater suppression of erastin2-induced cell death.

### Determination of GSH levels via Cayman GSH Kit

GSH levels in one experiment were determined using the Glutathione Assay Kit (Cayman Chemical, Cat. No. 703002). U-2 OS cells were seeded at a density of 100,000 cells/well in McCoy’s 5A medium in 6-well dishes. The next day, cells were washed once in warm HBSS, and the medium was replaced with DMEM minus Arg media, with indicated amino acid re-supplementation and small molecule treatment. After 16 hours, cells were washed once in ice cold PBS. Cells were scraped into 50 uL MES collection buffer and sonicated with 10 one-second pulses at maximum amplitude with a Fisher Scientific Model 120 Sonic Dismembrator (Thermo Fisher) to lyse cells. To exclude debris, lysate was centrifuged at 12,700 RPM for 15 minutes at 4°C and supernatant was isolated. 25 μL of lysate was aliquoted and stored frozen at −20 °C for under a week before a Bradford Assay was performed to determine the protein concentration for normalization. The rest of the lysate was deproteinated by adding one volume of .1 g/mL metaphosphoric acid (Acros Organics, Cat. No. 219221000), vortexing thoroughly, then centrifuging at 12,700 RPM for 3 minutes at room temperature. Metabolite extract was stored at −20 °C for under a week before being measured according to manufacturer’s instructions.

### Puromycylation assay

U-2 OS cells were seeded at a density of 100,000 cells/well in McCoy’s 5A medium in 6-well dishes. The next day, cells were washed once in warm HBSS, and the medium was replaced with DMEM-base resupplementation medium, as described above, with or without Arg, leucine, or INK 128 (1 µM). After 16 h, treatment media was aspirated and cells were pulse-labelled for 15 min in the same medium containing puromycin (10 μg/mL). At this point, cells were harvested and processed via Western Blot as described above. Primary antibody was mouse anti-puromycin (Clone 12D10, EMD Millipore, Cat No. MABE343) at a dilution of 1:5000 and rabbit anti-GAPDH at a dilution of 1:1000. LI-COR secondary antibodies were constituted in a 1:1 ratio of blocking solution and TBST at a dilution of 1:15,000. Membranes were imaged on the Odyssey CLx Imaging System (LI-COR Biotechnology, Lincoln, NE, USA). Results were quantified and normalized to 1 by computing the ratio of anti-puromycin signal to the GAPDH signal within each treatment condition. The results of three independent experiments were averaged and represented as mean ± SD.

### RT-qPCR

U-2 OS cells were seeded at a density of 100,000 cells per well in 2 mL of McCoy’s 5A medium in 6-well dishes. The next day, cells were washed once in HBSS then the medium was replaced with DMEM-base resupplementation medium, as described above, containing Arg or lacking Arg for 16 h. At this point, cells were carefully washed twice with ice cold HBSS. Cells were then gently scraped into 1 mL HBSS and centrifuged at 3,000 RPM for 2 min to pellet without disrupting the membrane. Supernatant was aspirated and RNA was extracted using the QiaShredder (Qiagen, 79654) and RNeasy (Qiagen, 74134) kits. Pellets were resuspended in 350 μL RLT buffer, then centrifuged through the QiaShredder column at 15,000 RPM for 2 min. 350 μL of 70% ethanol was added to the flow through and run through the RNeasy column for 15 sec at 15,000 RPM. Samples were then washed sequentially once with 700 μL RW1 and twice with 500 μL RPE before a one min spin to dry the column. RNA was eluted in 65 μL DNase/RNase free water and stored at −80°C until cDNA synthesis. cDNA synthesis reaction was performed using the Taqman Reverse Transcriptase Kit (Life Technologies, N8080234), with 5.5 mM MgCl_2_, 500 μM dNTPs, 2.5 μM DT oligos, 2.5 μM hexamers, 0.4 units/μL RNAse inhibitor and 3.125 units/μL reverse transcriptase. 1 μg of RNA was used per reaction. cDNA was synthesized using a ProFlex PCR System (Applied Biosystems, Foster City, CA) thermocycler with the following program: 25°C for 10 min, 48°C for 40 min, and 95°C for 5 min. cDNAs were stored at −20°C until qPCR reaction. qPCR reactions were run using SYBR Green Master Mix (Cat No. 4367659, Life Technologies) and run on the Applied Biosystems QuantStudio 3 real-time PCR machine (Thermo Fisher). Data were analyzed using the ΔΔCT method using *ACTB* as a control. Primers for qPCR were as follows: *ACTB* (F: ATCCGCCGCCCGTCCACA R: ACCATCACGCCCTGGTGCCT), *CBS* (F: TGAGATCCTGCAGCAGTGTG R:CTCCTTCAGCTTCCTGGCAA), *CTH* (F: CCAGCACTCGGGTTTTGAATA R: TGCCACTTGCCTGAAGTACC).

---

**Supplementary Figure 1.**
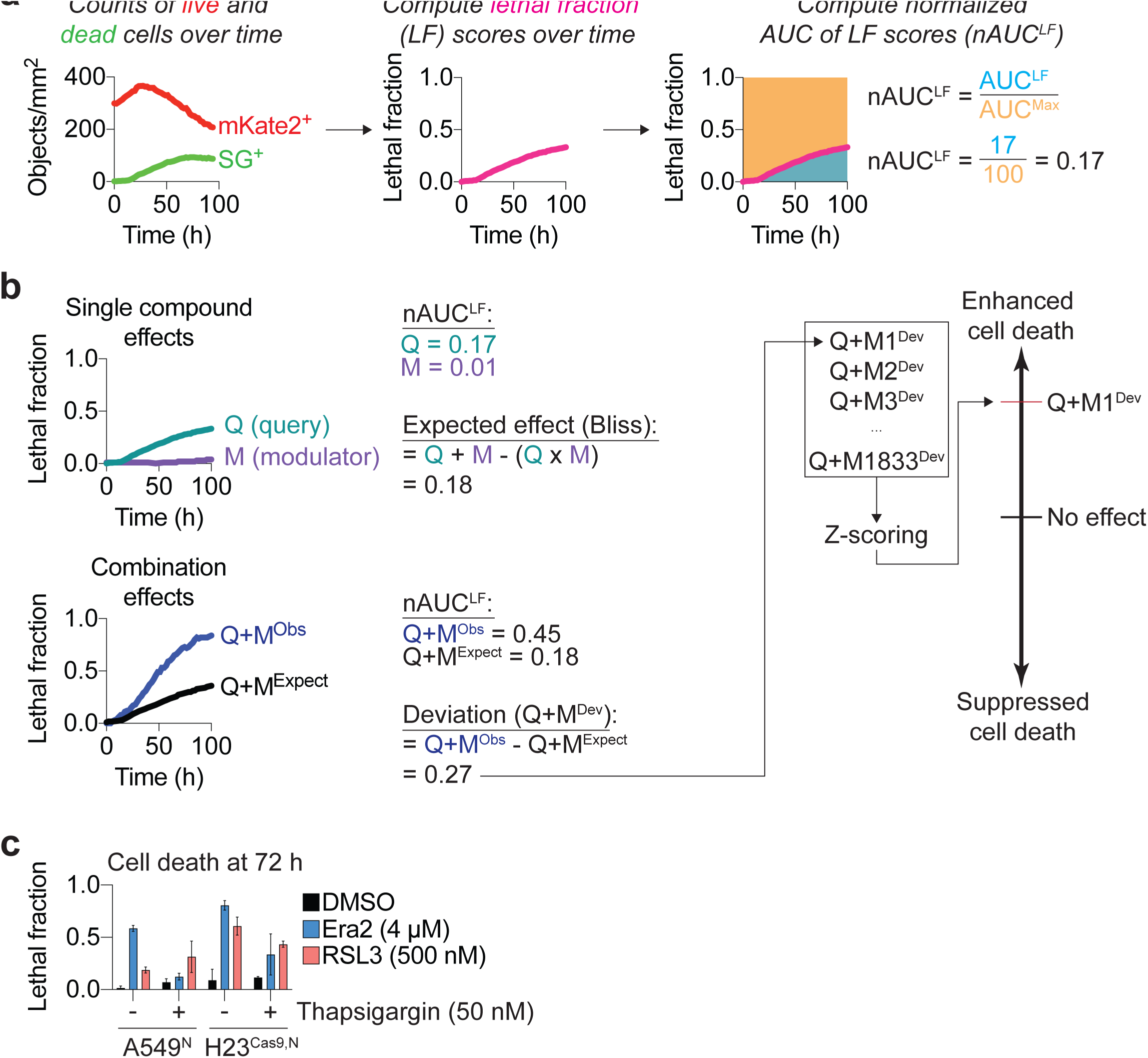
(a) Calculation of normalized area under the curve of lethal fraction (LF) scores (nAUC^LF^). (b) Calculation of modulatory effects using nAUC^LF^ scores. (c) Cell death in two different cell lines determined using the STACK methods in cells treated as indicated.

**Supplementary Figure 2:**
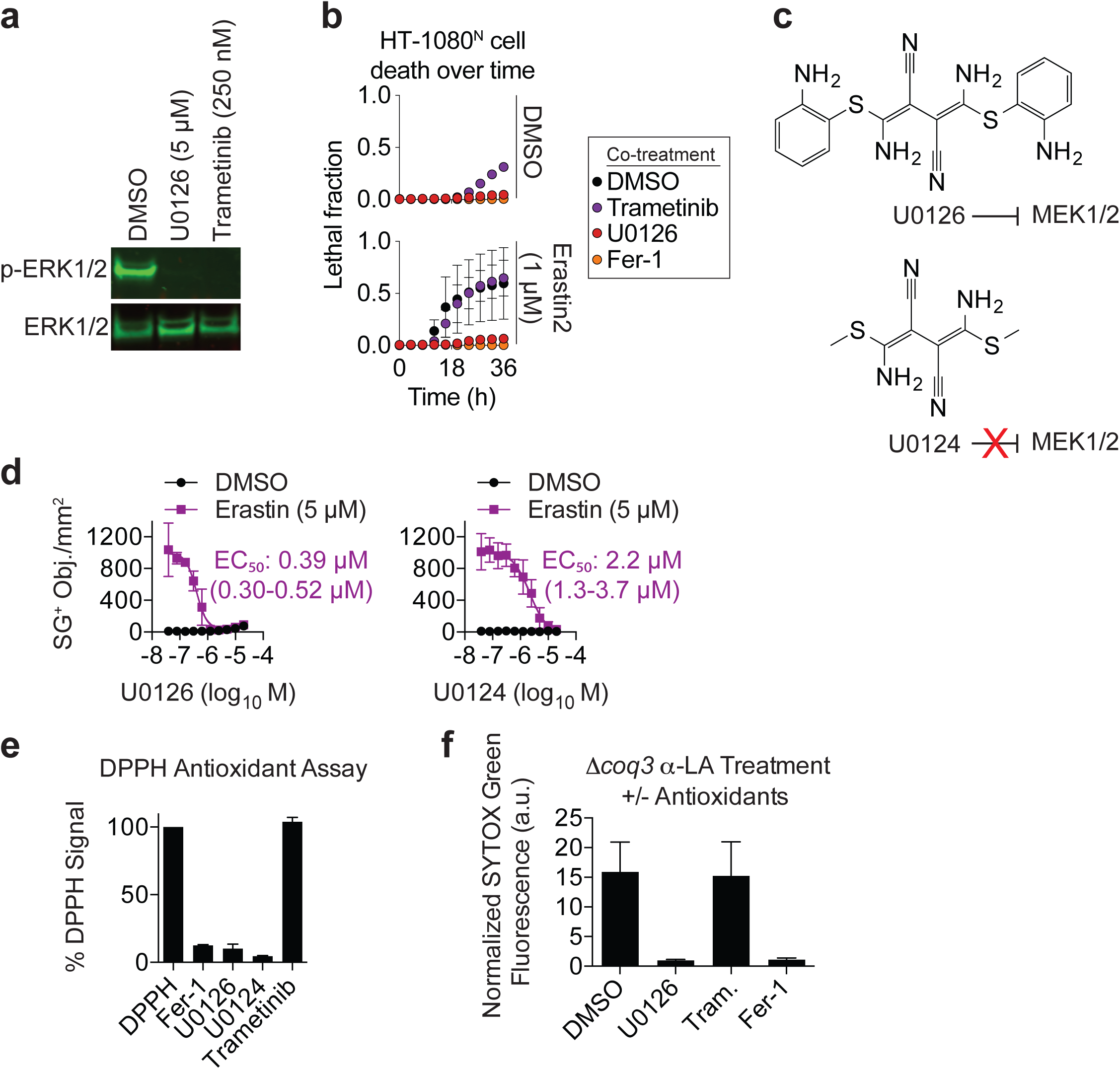
U0126 can act as an antioxidant suppressor of ferroptosis. (a) Western blot of serum starved HT-1080 cells stimulated with media containing 10% FBS for 30 min ± DMSO (vehicle), U0126 (5 μM) or trametinib (250 nM), an allosteric MEK1/2 inhibitor. Blots were probed for phospho-p44/42 MAPK (ERK1/2) and total MAPK (ERK1/2). One representative blot from three independent experiments is shown. (b) Cell death over time. Compound concentrations are as in (a), with ferrostatin-1 (Fer-1, 1 µM). (c) Structure of U0126 and the analog U0124. U0126 inhibits mitogen activated protein kinase kinase 1/2 (MEK1/2), while U0124 does not (Favata et al (1998) *J Biol Chem*). (d) SYOTX Green positive (SG^+^) dead cells at 24 h. Both U0126 and U0124 can inhibit erastin-induced ferroptosis, albeit with different potency. (e) Cell-free, free radical scavenging activity of U0126, U0124, and trametinib, monitored by changes in the absorbance of the stable free radical bearing molecule 2,2-diphenyl-1-picrylhydrazyl (DPPH) at 517 nm. Fer-1 = ferrostatin-1, a known radical-trapping antixodiant included as a positive control. (f) U0126 inhibits cell death in a yeast assay for polyunsaturated fatty acid autoxidation-induced cell death, while trametinib (Tram.) does not. α-LA: alpha-linolenic acid. All inhibitors were tested at 5 µM. Results in b,d, e and f represent mean ± SD from three independent experiments.

**Supplementary Figure 3.**
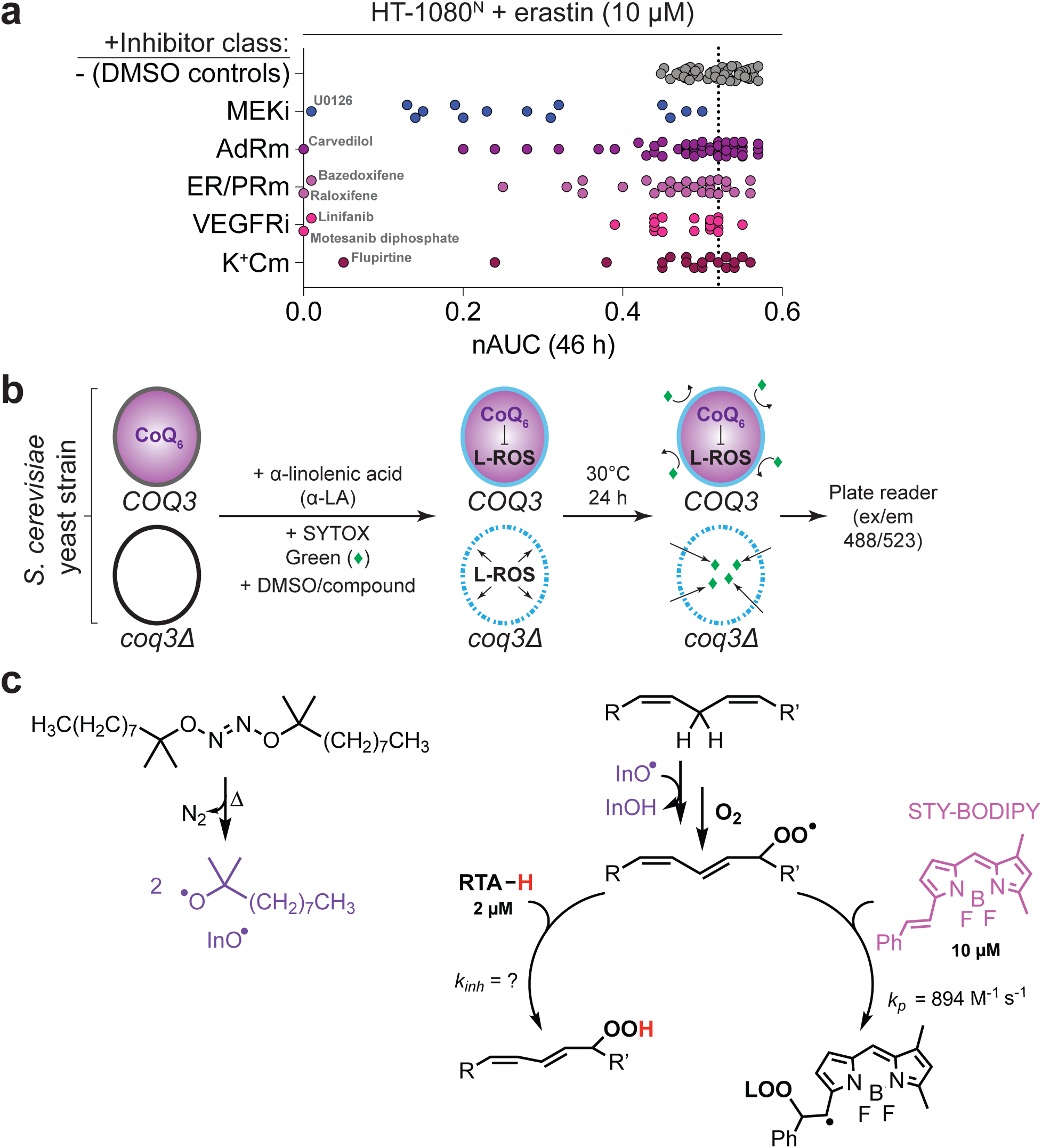
Investigating off-target ferroptosis inhibitors. (a) Data extracted from the profile presented in Fig. 1a for HT-1080^N^ cells treated with erastin (10 µM). Each dot represents a single compound, organized together by major target. All modulator compounds were tested at 5 µM. MEKi = mitogen activated protein kinase kinase 1/2 inhibitors (n = 14); AdRm: adrenergic receptor modulators (n = 55); ER/PRm: estrogen/progesterone receptor modulators(n = 29); VEGFRi: vascular endothelial growth factor receptor inhibitors (n = 20); K^+^Cm: potassium channel modulators (n = 19). The vertical dotted line indicates the average lethality for each lethal compound plus vehicle (DMSO) only. (b) Schematic of a *Saccharomyces cerevisiae (S. cerevisiae)*-based lipid autoxidation-induced cell death assay. α-linolenic acid (α-LA) is oxidized to form lipid reactive oxygen species (L-ROS). *COQ3* (i.e. wild-type) yeast produce Coenzyme Q_6_ (CoQ_6_) to detoxify L-ROS and prevent damage to the plasma membrane (solid blue line). CoQ_6_-defective *coq3Δ* yeast cannot detoxify L-ROS, resulting in a permeabilized (i.e., damaged) plasma membrane (dashed blue line). In the assay, *coq3Δ* yeast are co-treated with ethanol (EtOH) or α-LA (500 μM), the dead cell marker SYTOX Green (250 nM), and DMSO or the candidate antioxidant or iron chelator for 24 h. At the end of the incubation, cell death is measured by determining SYTOX Green uptake (excitation at 488 nm/emission at 523 nm). (c) Outline of the STY-BODIPY kinetic competition assay. Egg-phosphatidylcholine (1 mM) and STY-BODIPY (10 µM) are incubated with 0.2 mM di-tert-undecyl hyponitrite (DTUN), in addition to a radical trapping antioxidant (RTA-H).

**Supplementary Figure 4.**
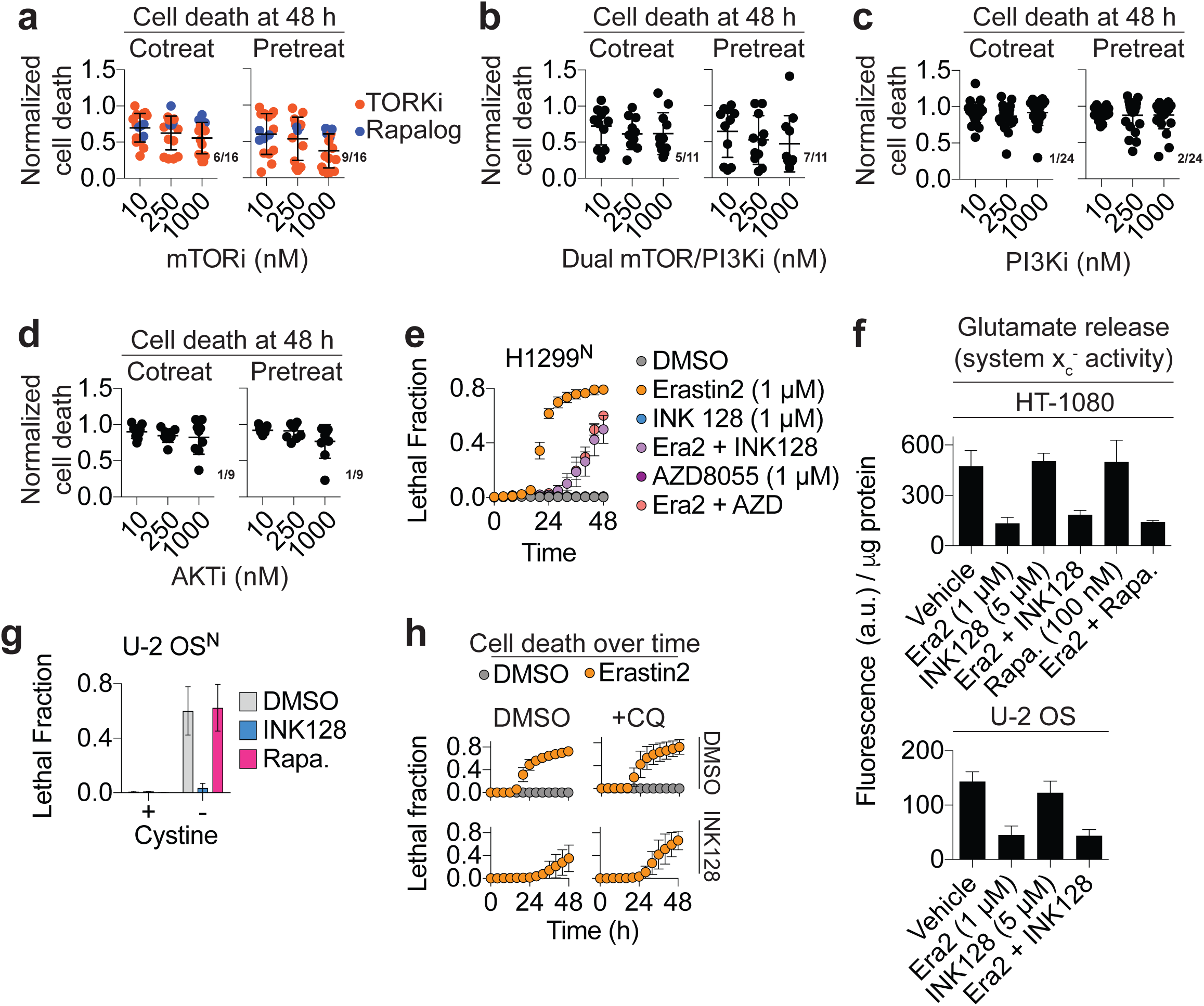
The role of mTOR signaling in ferroptosis. (a-d) Cell death of U-2 OS cells treated with erastin (20 µM) as determined by imaging of SYTOX Green positive dead cells at 48 h. Cell death for each condition is normalized to erastin-only controls (0 = alive, 1 = dead). Cells were either cotreated (Cotreat) or pretreated for 6 h (Pretreat) with small molecule inhibitors targeting mTOR (n = 16, a), mTOR/PI3K (n = 11, b), PI3K (n = 24, c) or AKT (n = 9, d). Each inhibitors was tested at 1000 nM, 250 nM or 10 nM. Each dot represents the effects of a single inhibitor in each class from one experiment. In (a), TORKis (n = 12) are colored black and rapalogs (n = 4) are blue. Numbers in pink represent the fraction of inhibitors in each class that suppress cell death by at least 50% at 1000 nM. (e) Cell death over time in H1299^N^ cells. (f) System x_c_^-^ activity inferred from glutamate release over 2 h from HT-1080 and U-2 OS cells treated as indicated. Era2: erastin2, Rapa: rapamycin. (g) Quantification of cell death at 48 h in U-2 OS^N^ cells in medium ± cystine ± INK128 (1 µM) or rapamycin (Rapa., 100 nM). (h) Cell death over time in U-2 OSN cells treated with erastin2 (2 µM) ± INK128 (1 µM) ± the lysosomal deacidifying agent chloroquine (25 µM). Results in e-h represent the mean ± SD from two to four independent experiments.

**Supplementary Figure 5.**
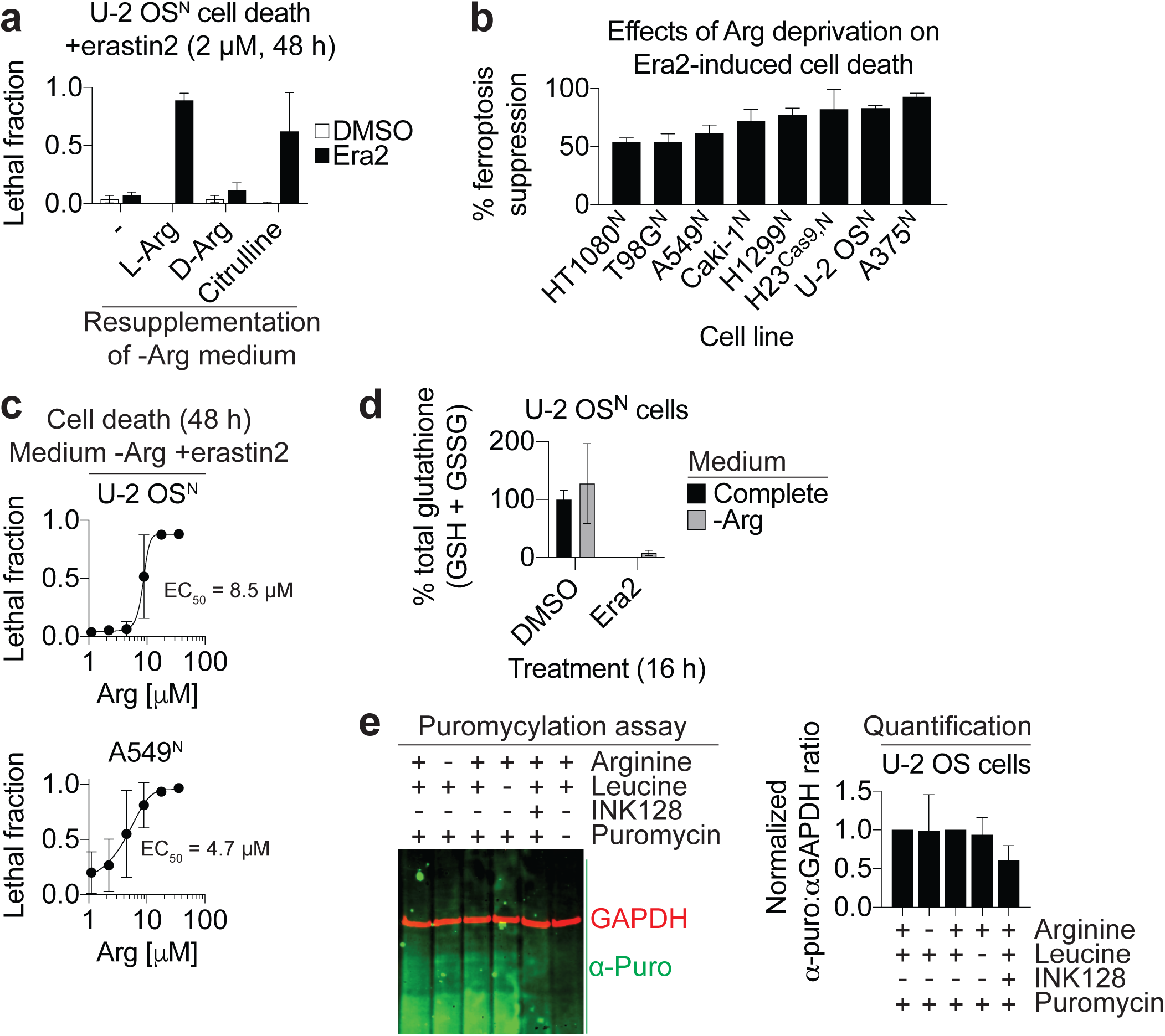
Arginine deprivation inhibits ferroptosis. (a) Cell death in -Arg medium resupplemented with L-arginine (L-Arg), D-arginine (D-Arg) or citrulline, each at 356 µM (the concentration of L-Arg in standard DMEM + dialyzed serum). (b) Percent ferroptosis suppression by growth in -Arg medium versus regular medium. Cell lines were grown in DMEM medium containing dialzed FBS ± Arg (356 µM) ± erastin2 (Era2) for 48 h. The concentration of Era2 was cell-type-dependent to achieve > 90% lethality + Arg, as follows: HT-1080^N^ (1 µM), T98G^N^ (2 µM), A549^N^ (4 µM), Caki-1^N^ (1 µM), H1299^N^ (2 µM), H23^N^-Cas9 (1 µM), U-2 OS^N^ (2 µM), A375^N^ (2 µM). The formula for computing % ferroptosis suppression is given in the Methods. Note: values closer to 100% indicate greater suppression of erastin2-induced cell death (i.e. less cell death). (c) Dose-dependent effect of Arg resupplementation on erastin2-induced cell death. Erastin2 concentration was the same as in b. (d) Total glutathione (GSH + GSSG) expressed as a percentage of the DMSO control grown in complete medium. Erastin2 (Era2) was used at 2 µM. (e) Detection of puromycylated peptides. U-2 OS cells were incubated as indicated (note: INK128 was used at 1 µM) for 16 h then pulsed for 15 min with puromycin, prior to cell lysis. Results in b-e represent the mean ± SD from at least three independent experiments.

